# Abundant positively-charged proteins underlie JCVI-Syn3A’s expanded nucleoid and ribosome distribution

**DOI:** 10.1101/2025.08.16.670686

**Authors:** Gesse Roure, Vishal S. Sivasankar, Roseanna N. Zia

**Affiliations:** Department of Mechanical and Aerospace Engineering, University of Missouri Columbia, Missouri, United States of America

## Abstract

Nucleoid compaction in bacterial cells has been attributed to cytoplasmic crowding, supercoiling effects, and the action of nucleoid-associated proteins (NAPs). In most bacteria, including *E. coli*, these mechanisms condense the nucleoid to a smaller volume within the cell, excluding most ribosomes to the surrounding cytoplasm. In contrast, the nucleoid in many *Mycoplasma*s, including the *Mycoplasma*-derived synthetic cell JCVI-Syn3A, spans the entire cell, with ribosomes distributed throughout. Recent models of Syn3A representing only DNA and ribosomes (both charge neutral) instantiated the experimentally-observed expanded nucleoid and ribosome distribution. However, we found that this configuration becomes dynamically unstable, giving way to a compacted nucleoid that expels ribosomes to the periphery, suggesting the need for a more detailed model. Speculation emerging from recent studies of Syn3A suggests that its lower concentration of NAPs underlie its expanded nucleoid. We are interested in this genotype-to-’physiotype’-to-phenotype implication: that coupled transcription, translation, and nucleoid remodeling lead to different phenotypical outcomes. We developed a coarse-grained computational model of Syn3A, physically and explicitly representing ribosomes, cytoplasmic proteins, and a sequence-accurate chromosome with physiological distributions of size, charge, and relative molecular abundance. An interplay between Brownian dynamics, DNA stiffness (both inherent and NAP-enhanced), and electrostatic charge led naturally to a stable molecular distribution. We find that an interplay between inherent and induced DNA stiffness, heterogeneous mesh size, and crowding enhances nucleoid compaction and ribosome expulsion via a competition between entropic and enthalpic forces. In contrast, electrostatic interactions and size-polydispersity counteract these effects and expand the nucleoid. In particular, Syn3A’s atypically high abundance of positively-charged proteins shields ribosomes’ negative charge, allowing them to interpenetrate the nucleoid. Finally, we observe condensate formation arising from electrostatic interactions with potential implications on transcription and translation rates.

**Author summary:** The chemical elements of DNA in cells — its genes — carry the instructions for it to grow, replicate, and divide but its physical organization keeps those instructions in order and easy to read. In bacterial cells, DNA occupies a specific region that determines where its mRNA transcripts are ultimately translated by ribosomes into the proteins that help the cell grow and adapt. This structure differs markedly between bacteria in two prominent genera: *Escherichia* and *Mycoplasmas*. Previous models and experiments explain why *Escherichia*’s DNA tends to be compact and centralized, expelling ribosomes to the cell periphery, separating transcription from translation. But many *Mycoplasmas*, much simpler cells, have expanded DNA — and their ribosomes are co-located within the DNA’s structure. It’s been proposed that the same proteins that help cells adapt to stress may underlie this behavior, and they are far less abundant in *Mycoplasmas*, suggesting that expanded DNA may help improve low adaptability. To interrogate the interplay between mesoscale physical DNA architecture, stress proteins, and physical interactions in the cell, we built a computational model of the *Mycoplasma*-derived minimal cell JCVI-Syn3A, representing all of its DNA, ribosomes, and proteins with explicit physical resolution and interactions. By testing the conditions that drive these divergent “physiotypes”, we found an explanation in Syn3A’s atypically positively-charged proteins, which cloak ribosomes, letting them enter and expand the nucleoid.

## Introduction

The chromosomal DNA in most bacteria is a long, closed-loop molecule that packs densely within the cell but, unlike eukaryotes, is not enclosed by a nuclear membrane. Instead, its mesh-like structure forms its own well-defined region. This region, called the nucleoid, contains the majority of the chromosomal DNA, as well as a host of molecules involved in gene expression within the solvent-filled ‘pores’ of the mesh network. The physical features of bacterial DNA span multiple length scales, ranging from localized bends over a few base pairs, to supercoils, to macrodomains. The structure at each of these length scales is associated with functional interactions with transcription factors and other molecules, which together regulate gene expression. Its longest length scale is its overall or ‘global’ size. In most bacteria, including *Escherichia coli*, the global envelope of the nucleoid is smaller than the whole cell, leaving the nucleoid surrounded by chromosomal DNA-free cytoplasm. In such cells, the ribosomes are almost entirely excluded from the nucleoid region (besides small subunits) [1–4]. This physical exclusion separates gene transcription within the nucleoid from mRNA translation outside it [5]. This segregation of macromolecules, and its corresponding spatial organization, produces spatially heterogeneous transport rates, which can in turn affect reaction rates [5–7]. For example, one consequence is the sequestration of macromolecules that helps speed mRNA translation [8–10]. The volume of this compact nucleoid region can change with growth rate and other conditions, which may, in turn, change the packing density within the nucleoid [2]. As another example, gene expression rates are partly regulated by changes in DNA structure, ranging from local bends to supercoils to global compactness, all of which can physically hinder or enhance access to genes. We call these links between physical structure and phenotype, “physiotype-to-phenotype” connections. Previous experiments and models suggest that compaction and expansion of the nucleoid is driven partly by crowding [11], as well as by nucleoid-associated proteins (NAPs) [12], a class of DNA-binding proteins homologous to the histone proteins found in eukaryotic cells. NAPs serve a range of functions by binding to DNA and generating mechanical forces that can locally remodel DNA by creating bends [13], making it stiffer [13], and/or bridging separated gene segments together [14, 15]. Some of these local effects, such as the bridges generated by NAPs such as H-NS, are known to influence nucleoid compaction, as observed both experimentally [16] and in computational simulations [17]. Although some contributors to DNA remodeling have been studied to varying extents individually, there are still important open questions, including why some bacterial nucleoids span the entire cell.

In contrast to most bacteria, the nucleoid in some *Mycoplasmas* spans the entire cell [18], with ribosomes distributed throughout — including within the nucleoid [19].

Related to this expanded structure, *Mycoplasmas* have far fewer NAPs [20, 21] and, given their known role in nucleoid compaction, this paucity of NAPs suggest one mechanistic explanation for the expanded nucleoid. However, other cytoplasmic conditions such as crowding, depletion interactions, and electrostatics, that are well known to compact DNA [22] may also undergird this behavior, motivating further exploration. Mechanistic understanding of how DNA interacts with cytoplasm to undergo *physical* restructuring may also uncover new insights about how the nucleoid orchestrates cell-wide dynamics to regulate transcription and translation during growth and during adaptation to environmental changes. Computational modeling provides a useful complement to experiments, owing to the difficulty of dynamically imaging restructuring *in vivo*. In this work, we develop a new computational model of a representative *Mycoplasma* cell to mimic and shed light on the physico-chemical interplay between DNA and cytoplasm that produces an expanded nucleoid.

Models of DNA’s multi-scale structure, condensation, and remodeling include thermodynamic theory, polymeric bead-spring models immersed in mean-field backgrounds, and detailed computational models of DNA interacting with explicitly simulated cytoplasm. The primary difference between these three approaches is the degree of physical resolution of DNA interaction with itself and its surroundings. We briefly review these approaches, each of which contributes to current understanding of nucleoid structure.

Thermodynamics-based approaches model restructuring of DNA using two primary frameworks: polymer collapse and phase separation. De Genne’s fundamental works on polymer collapse suggested that solvent quality may play an important role in DNA condensation [23], setting the foundation for many subsequent thermodynamic models. As another example, Post and Zimm’s expanded thermodynamic theory modeled three different coiling motifs as phases, producing an early mechanistic concept of how the bacterial nucleoid initially forms [24]. De Vries later expanded de Genne’s solvent quality framework, adding theoretical effects of cytoplasmic crowders [25]. Building upon the idea of phases, Odijk used a liquid-like phase-separation model to put forth the idea that bacterial DNA supercoils condense together due to depletion interactions between them induced by cytoplasmic crowders, offering another route to nucleoid formation [26]. These studies verified experimental observations of the compacting action of protein crowders [11]. Their low (or zero) computational cost makes these models highly desirable, driving ongoing expansion of the theory. They are also frequently used to validate computational models [27–29]. However, these thermodynamics-based frameworks smear out physical resolution, losing *local* spatial conditions, such as crowding and explicit DNA interactions that can suppress or silence genes.

Dynamical models of DNA with physical resolution range from all-atom computational frameworks focusing on local dynamics to continuum models that combine theory and simulations to explore system-level behaviors. For physical structure and dynamics at the smallest scales, near-atomistic and all-atom modeling reveals changes over corresponding ranges in DNA and other cytoplasmic biomolecules [30–33]. For example, the binding interactions between NAPs and DNA that produce local bending and form bridges, and the condensing action of amino acids, ions, and polypeptides, can be modeled and studied with corresponding detail using atomistic simulations [34–37]. Combined with AI tools such as AlphaFold [38], atomistic modeling has revolutionized structural biology — revealing, for example, the complex molecular-level dynamics of protein folding [39–41] and how binding interactions change local DNA conformation [42]. However, protein-folding simulations still require intermediate coarse-graining steps, and the studies of DNA binding interactions are limited to partial DNA segments of fewer than 100 base pairs, owing to the steep computational cost per atom in simulation [43]. Overall, to simulate larger or more complex molecules and handle hundreds of thousands of base pairs, atomistic simulations must be combined with coarse-graining approaches [44–46]. Predicting cooperative and global deformation of the bacterial nucleoid is thus well beyond the capability of all-atom models alone.

To investigate such global DNA dynamics, continuum theories have been developed to model macroscopic behaviors away from equilibrium. Examples include field-theorical diffusion [47] and active-fluid models of chromatin transport [48]. This comprehensive, system-wide dynamical modeling of DNA probes far-from-equilibrium emergent behaviors not captured by prior thermodynamics theory, predicting how competition between coiling and diffusion swells the entire bacterial nucleoid [47], and how fluid flow inside the eukaryotic nucleus drives coalescence of DNA [48, 49]. The physical resolution sacrificed by continuum models is an excellent tradeoff for these discoveries. However, even though they encompass the entire nucleoid, the smearing-out of detailed DNA structure means they are not amenable for connecting atomistic rearrangements following binding interactions to both local and global nucleoid remodeling.

Mesoscale biophysical models hold promise for revealing mechanistic connections between atomistic and macroscopic length scales, which is crucial for understanding how DNA mediates cell-wide function. For example, coarse-grained polymer models represent macromolecules as a chain of mesoscale, colloidal particles. These bead-spring models [50] have been widely used to capture global behavior of large macromolecules, including the global dynamics of DNA.

Mesoscale, physically resolved DNA models permit study of multi-scale DNA remodeling and, when modeled within cytoplasm, can reveal how DNA structure affects cell-wide biomolecule organization and transport. Bead-spring polymer models predict DNA’s local dynamics and how these lead to multi-scale structure [51, 52], explicitly modeling the dynamics of supercoiling [53–55], knot dynamics [56], replication [57], condensation [58], and macromolecular transport within phase-separated domains [59]. The ability to model these large-scale phenomena with a sufficiently detailed description of local DNA interactions makes these models very suitable for study of chromosome-level DNA remodeling and its effects on nucleoid compaction and cytoplasm organization. As an early example, Sottas et al. [54] used coarse-grained simulations to show that dynamical supercoiling and cytoplasmic salt concentration affects compaction of plasmid DNA. Joyeux’s recent Brownian simulations of a generic bead-spring model nucleoid showed that confinement and crowding can drive nucleoid compaction even in the absence of supercoiling [60]. Later model improvements accounted for supercoiling and DNA-bridging proteins, which were shown to induce further compaction [17, 61]. Other simulations have reinforced these observations, and introduced additional effects such as crowder-size bidispersity and non-spherical confinement [3, 4, 62]. In contrast to thermodynamic theories, these approaches explicitly model local physical interactions from crowders and bridging proteins. A recent combined experimental and mesoscale simulation study of cytoplasm organization in *E. coli* revealed how nucleoid structure and charge affects macromolecular diffusive transport and segregation of large macromolecules [7]. Overall, these physically-resolved, coarse-grained approaches elucidate nucleoid restructuring as a response to interactions with simple cytoplasm or, separately, biomolecule organization driven by nucleoid features.

Despite this progress, why the nucleoid in some bacteria is always relatively compact while in other bacteria, it expands to fill the entire cell, is unknown. A compacted or expanded condition obviously correlates with distinctly different cell-wide biomolecule spatial distribution, but whether this organization drives compaction or vice versa is also unknown. Finally, two additional key drivers may further contribute to the compaction / spatial distribution condition: electrostatic charge and size polydispersity, both of which are implicated in condensate formation and segregation [63, 64]. Recent physically-resolved whole-cell models of *Mycoplasma*-derived synthetic cells introduced 10 base-pair resolution models of the complete DNA in a simulated simple cytoplasm of cytosol and ribosomes recapitulated DNA replication [57], with others also including RNA polymerase to visualize cell organization [65]. Both studies neglect all other cytoplasmic proteins, electrostatic charge, and nucleoid-associated HU proteins. Both studies also placed ribosomes in the experimentally measured cell distribution. In the former the DNA was self-assembled in the remaining volume; in the latter, the RNAPs were then added, followed by self-assembly of the DNA. Because the space was available during simulated self-assembly, a fully expanded nucleoid resulted. But the physiological distribution of other cytoplasmic proteins would tend to compact the nucleoid, so including this physiological contribution, along with electrostatic charges, will surely reveal more.

In this study, we examine several factors that contribute to the expanded nucleoid of *Mycoplasma*-type cells, including factors such as charges and DNA-bending HU proteins, which have been neglected in past models. To do so, we use a mesoscale-resolution whole-cell model of the synthetic cell JCVI Syn3A, a bacterium derived from *M. mycoides*, engineered to have the minimal genome capable of life [66, 67]. The simplicity of Syn3A’s genome and proteome is conducive to a whole-cell model and, because its genome is restricted to life-essential genes and has broad homology to all *Mycoplasmas*, several conclusions drawn from Syn3A be broadly applied to all *Mycoplasmas* [66], giving broad understanding regarding their nucleiods’ characteristics. Interest in Syn3A as a model bacterial cell has prompted its experimental study [68–70] and several models, ranging from kinetics-based whole cell models [19, 71] to nearly-all atom models [72] providing a rich basis for comparison. A recent review compares the benefits of this range of whole-cell modeling approaches [73].

## Results and discussion

### Recapitulating experiment-matched models

As a starting point, we build a baseline model for JCVI Syn3A and compare our molecular distribution to experimental results. To this end, we use the ribosome data reported by [19], which has been previously used to inform other Syn3A models [57, 65]. In the works by [57] and [65], a model Syn3A cell was built by placing ribosomes according to experiments, holding them fixed, and letting the DNA assemble itself around it. By using our swelling Monte Carlo algorithm, we generate an initial condition through this protocol, obtaining a cell configuration with the same almost-uniform ribosome distribution and a cell-spanning nucleoid, as seen in **Figure 1(a)**. Similarly to [57], the system shown in **Figure 1** comprises DNA, ribosomes, and entropic interactions between them (see *Methods* for more details). By standard, this approach recovers the experimental ribosomal distribution for the initial state. However, it is not clear if such configuration is dynamically stable, as entropic interactions are known to induce ribosome segregation and nucleoid compaction [60]. To investigate the stability of this macromolecular configuration, we allowed Brownian motion to commence, resulting in the configuration shown in **Figure 1(b)**. Our results indicate that the entropic interactions between ribosomes and DNA beads led to substantial — but not total — migration of ribosomes toward the cell membrane, resulting in an accumulation of ribosomes near the membrane and a moderate compaction of the nucleoid, indicated by a DNA-poor region near the membrane and higher DNA density near the cell center. Alternatively, our algorithm allows us to generate the configuration of ribosomes and DNA simultaneously without *a priori* knowledge of the experimentally-measured ribosome configuration. The resulting configuration using this approach is shown in **Figure 1(c)**. The configuration in **Figure 1(c)** is strikingly similar to the one obtained by dynamically evolving the initial conditions from experimental measurements [**Figure 1(b)**], and also shows an accumulation of ribosomes near the cell membrane and a slightly compacted nucleoid. Overall, the results in **Figure 1** show that, in the presence of entropic exclusion, ribosomes diffuse naturally out of the nucleoid and accumulate near the cell membrane, compacting the nucleoid by at least 15% by volume. Such nucleoid compaction and ribosome exclusion are inconsistent with experimental observations of near-homogeneous ribosome distribution in Syn3A, indicating that such a simple model (only ribosomes and DNA; purely hard-sphere repulsive interactions [57]) does not sufficiently capture the physics required to recover the experimentally measured macromolecular distribution. Indeed, in addition to DNA and ribosomes, Syn3A cells have a multitude of biomolecules of widely varying size, charge, and other characteristics that influence their dynamics and interactions. We expect this molecular composition, and corresponding interactome, to influence overall distribution of macromolecules and DNA. Beyond Syn3A’s well-defined proteomic size, charge distribution and packing fraction, it was recently proposed that the low count of nucleoid-associated proteins may also be a substantial driver of nucleoid compaction or expansion [74]. We expect that physiologically accurate combinations of these factors will improve our model’s prediction of the DNA and ribosome distribution reported in experiments — furthering the understanding of why many bacterial nucleoids are compacted while others are not. In the next several sections, we systematically investigate the following material properties of DNA and interactions on macromolecular distribution: ‘inherent’ or global DNA stiffness; local DNA bending produced by nucleoid-associated proteins; crowding; electrostatic charge, and how each of these effects influence macromolecular distribution in our model Syn3A.

**Fig 1.**
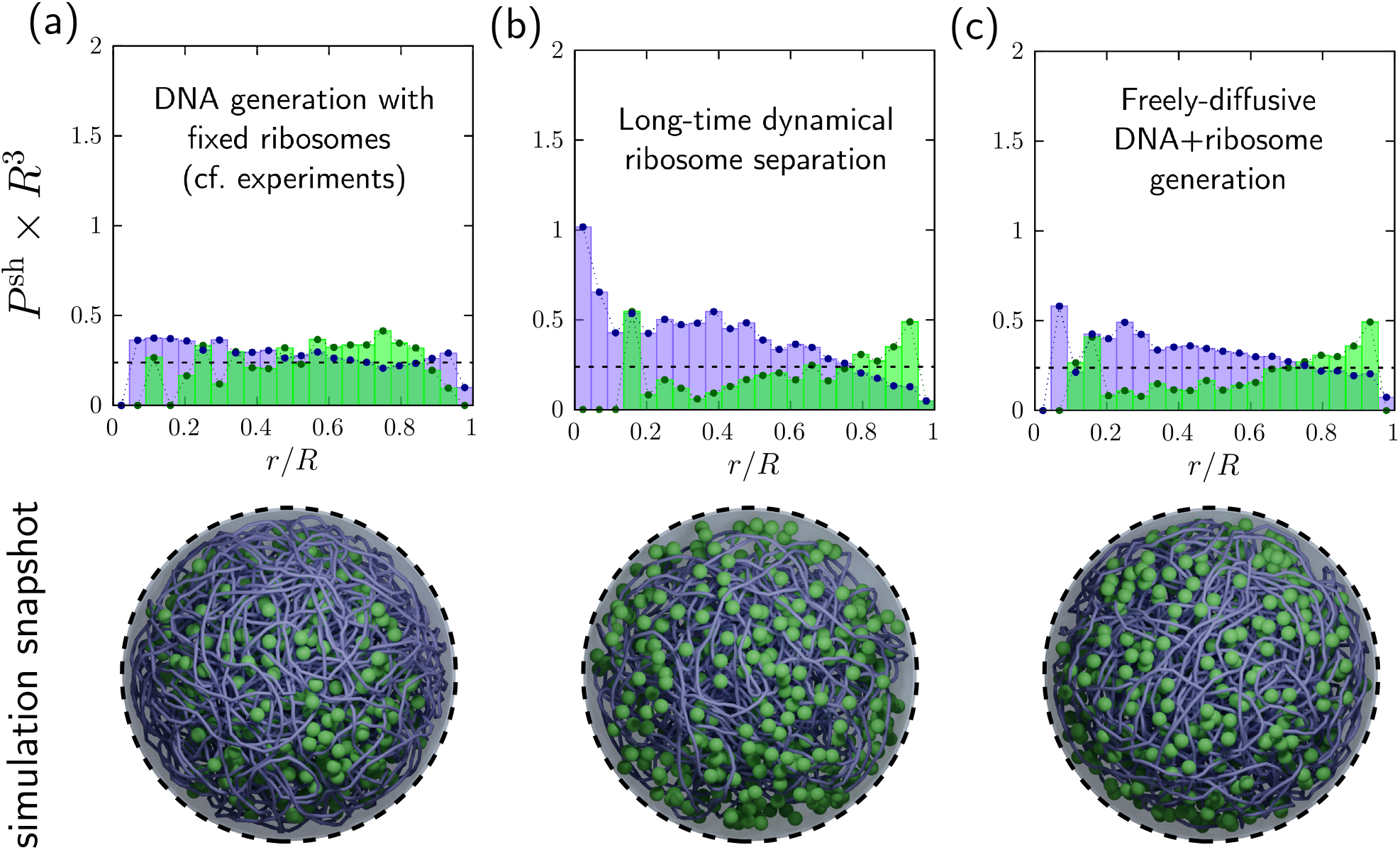
Nucleoid and ribosome spatial configurations predicted by our simplest computational model of JCVI-Syn3A, with only DNA and ribosomes (both charge neutral) present. Top row: Probability per unit volume of finding a DNA bead (purple) or ribosome (green), binned into annular shells from center (*r/R* = 0) to cell membrane (*r/R* = 1) in simulations. (a) Initial condition (no dynamics) set by experimental data from [19]. Both the DNA and ribosomes are homogeneously distributed throughout the cell. (b) Same initial condition as (a), but after the system has evolved dynamically via Brownian motion and entropic exclusion. Now the DNA has compacted toward the center, expelling ribosomes to the periphery. This show that dynamic modeling Syn3A with only charge-neutral DNA and ribosomes produces a configuration that does not agree with experiments. (c) New initial condition: DNA beads and ribosomes are initially distributed uniformly throughout the cell, after which the DNA beads self-assemble into a single continuous loop (nucleoid) via the swelling Monte Carlo algorithm [see *Methods*]. The model recovered the same prediction of compaction and ribosome exclusion without using experimental data, showing robustness of the self-assembly method, and reinforces that modeling DNA and ribosomes alone is insufficient to recover experimental measurement of homogeneous distribution. Bottom row: Simulation snapshots with 503 ribosomes and sequence-accurate nucleoid, coarse-grained to 100 base pairs per bead (smoothed for visual appeal).

### Nucleoid stiffness can induce DNA compaction

Base-pair stacking, controlled by chemical bonds, imparts DNA’s intrinsic multi-scale structure and gives rise to its mesoscale material properties such as bending, stretching, and torsional stiffness [75, 76]. As with non-biological polymers, we expect these intrinsic material properties to influence the nucleoid’s overall compactness and size. More generally, the chromosome includes both DNA and bound proteins that can further alter the nucleoid’s local and global stiffness, described in our model via dimensionless scalar coefficients *σ*_*s*_ and *σ*_*b*_, which set the chromosome’s resistance to stretching and bending, respectively [see *Methods*, Equations (4) and (5), and **Figure 8**]. In our baseline model, only the DNA’s base-pair stacking influences its bending stiffness (no binding proteins yet). That model (and corresponding results in **Figure 1**) has a uniform bending stiffness coefficient *σ*_*b*_ = 10, which is consistent with established literature values for ‘bare DNA’ (i.e., no nucleoid-associated proteins), obtained both by theoretical predictions and experimental measurements [76]. Similar values have also been used in recent coarse-grained computational models with simplified cytoplasmic constituents [17, 77] as well as Syn3A [57]. An intrinsically stiffer (or less-stiff) nucleoid could result in global expansion or compaction. To investigate how stiffening affects compaction, we simulated a range of values of *σ*_*b*_ and measured the impact on ribosome distribution, with a view toward improving agreement with experimental measurements. **Figure 2(a)** shows simulation snapshots for three different values of bending stiffness representing a soft nucleoid (*σ*_*b*_ = 0), a moderately stiff nucleoid (*σ*_*b*_ = 10) representing physiological conditions of Syn3A, and a stiff nucleoid (*σ*_*b*_ = 50). To obtain adequate statistics and minimize influence of DNA knotting and dynamical accessibility (e.g., kinetic trapping), we average the data across 100 simulations, and calculate the ensemble-averaged spatial distribution of DNA beads and ribosomes [see *Methods*].

**Fig 2.**
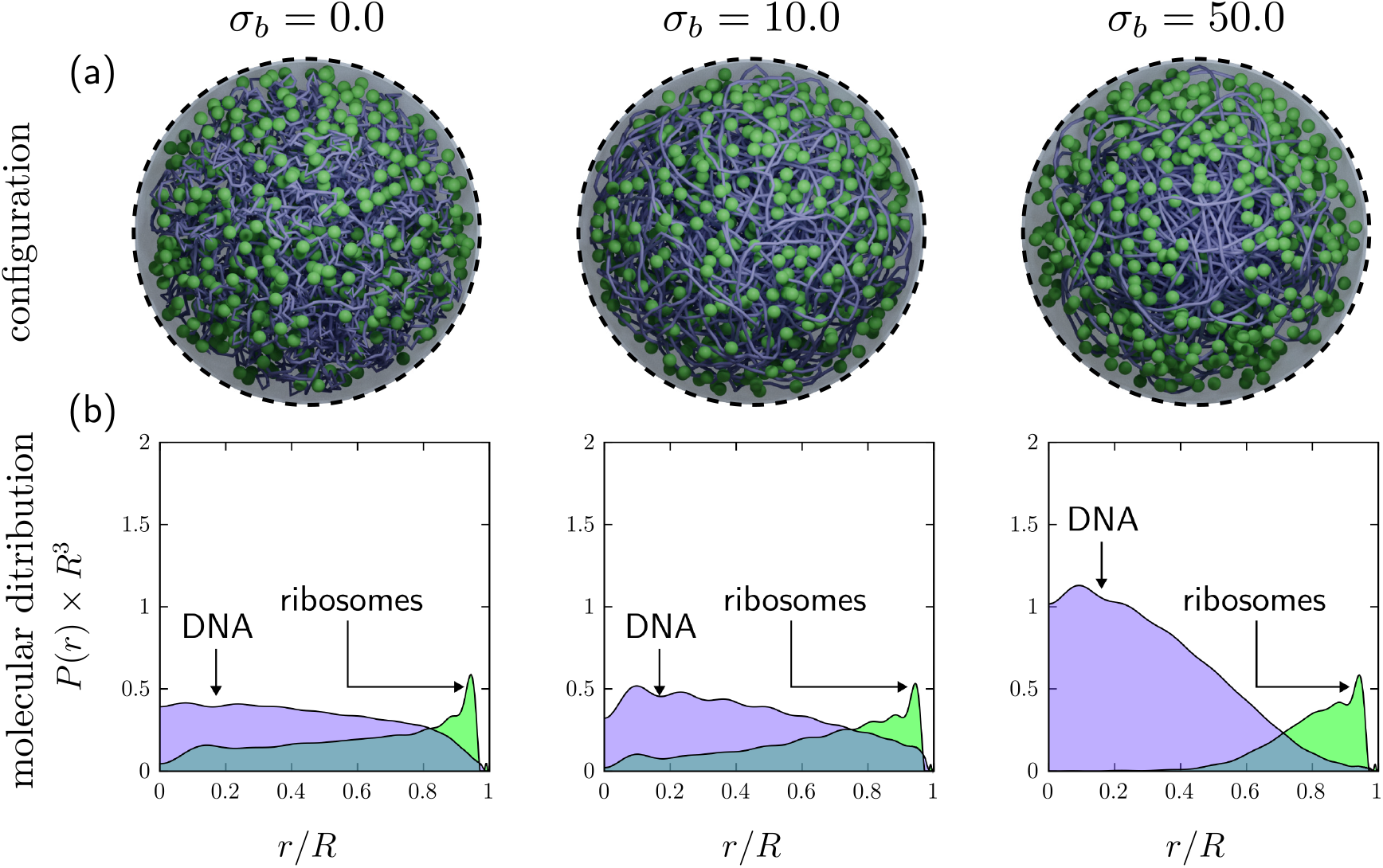
Impact of inherent nucleoid stiffness on distribution of DNA and ribosomes. Stiffness, *σ*_*b*_ (shown at top) is homogeneously distributed across all triplets in the DNA. (a) Simulation snapshots. Ribosomes: green spheres. DNA: purple nucleoid, 100-bp per bead (smoothed for visual appeal). (b) Plots of spatial distribution of DNA beads (purple) and ribosomes (green), as it varies from cell center to membrane. Data averaged over 100 independent dynamics simulations. No experimental data for multiple measurements are available.

**Figure 2(b)** shows that intrinsic stiffness alters both DNA compactness and ribosome distribution. An intrinsically stiffer nucleoid is more compact than a softer one, and this stiffness promotes ribosome exclusion: the stiffer the nucleoid, the more ribosomes are pushed away from the central portion of the cell. However, even the softest nucleoid (i.e., *σ*_*b*_ = 0) excludes some of the ribosomes into a DNA-poor region.

Overall, no value of inherent DNA stiffness recovers the experimentally measured distribution of ribosomes in the simplified models used here and previously [19, 57]. We also note that the changes in bending stiffness from *σ*_*b*_ = 10 to *σ*_*b*_ = 0 only weakly affects macromolecular distribution. In contrast, a stiffness increase to *σ*_*b*_ = 50 yields a nearly DNA-free region near the membrane that is occupied only by ribosomes; ribosomes are also almost completely excluded from the central portion of the cell. Note that these results pertain to a quite simplified model, with no electrostatic charge, no proteomic crowding, and none of the nucleoid-associated proteins (such as HU) known to induce changes in bending stiffness both *in vivo* and *in vitro* [78]. Before we explore how these additional effects influence molecular distribution, we first investigate the interplay between DNA stiffness, its physical structure, and the ribosome distributions observed in **Figure 2**.

### Nucleoid pore distribution partly explains ribosome exclusion

We speculate that the nucleoid’s inner pore-size or mesh-size distribution [80] at least partly explains why most ribosomes cannot reside within the nucleoid. To characterize the porous microstructure of the nucleoid, we carry out a Voronoi-based analysis following the approach by [79], with modifications that account for membrane confinement. Voronoi tesselation reveals pores of a range of sizes in the void regions excluded by DNA and confinement **Figure [3(a)(i)]**. We omit Voronoi edges near the membrane that produce spuriously high data for voids of nearly-null size. The Voronoi network is shown in green in **Figure 3(a)(ii)**. Each individual void is a segment of a network path, and is represented by a network edge *E*_*i*_ that is the center of a cylinder of diameter *d*_*i*_, where *i* = 1…*K* and *K* is the total number of nucleoid pores [**Figure 3(a)(iii)**]. This diameter sets the size of the largest particle that can fit within (pass through) the pore (segment).

**Fig 3.**
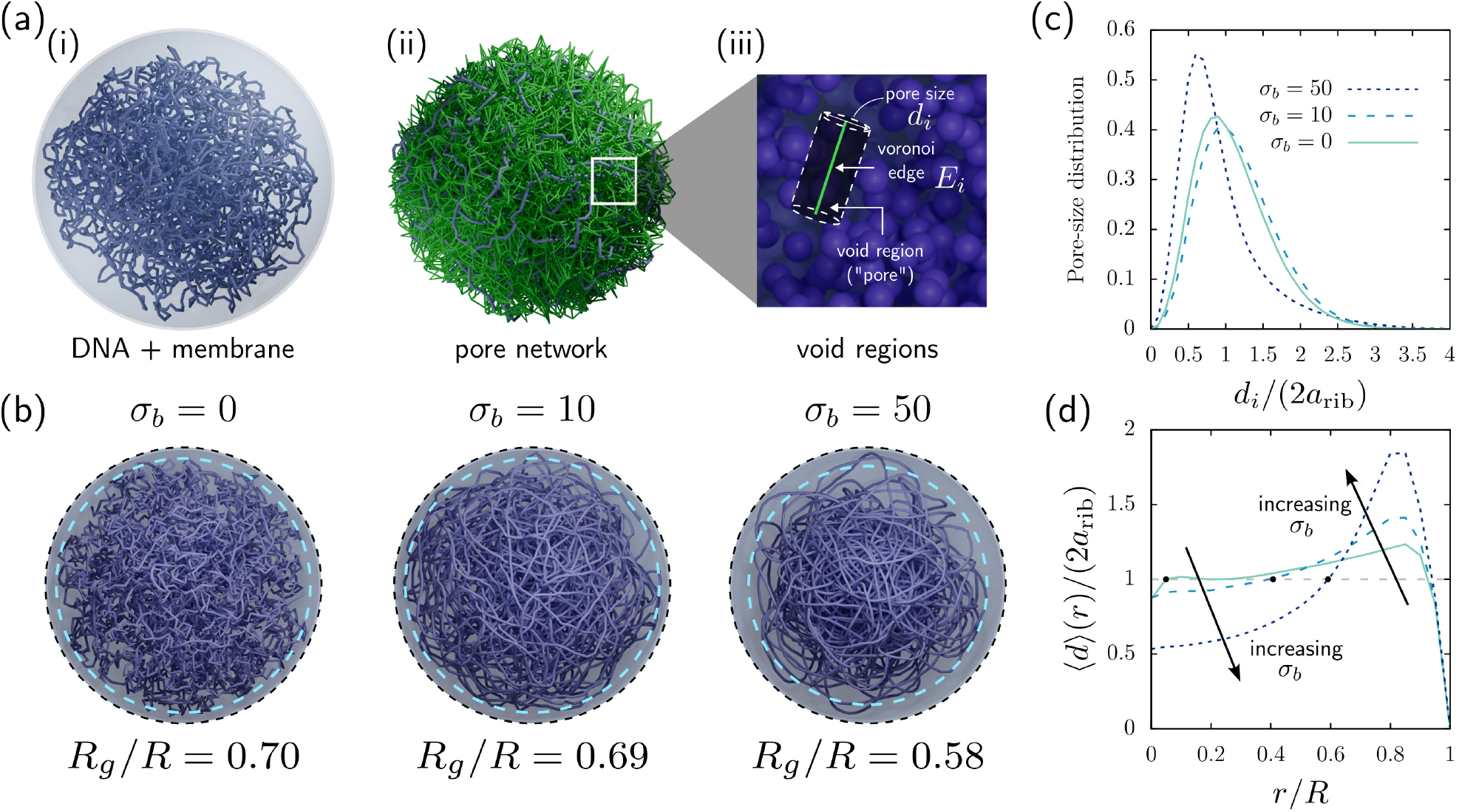
Voronoi analysis of nucleoid pores (mesh size) for three values of bending stiffness, as indicated in legends. (a) Voronoi interrogation of the nucleoid. (i) Simulation image showing only the DNA and enclosing membrane (ribosomes made invisible for visual clarity). (ii) Voronoi tesselation throughout the nucleoid. The resulting (infinitesimally thin) edges, shown in green, traverse the interconnected voids within the nucleoid. (iii) Determination of the radius *d*_*i*_ (pore size) of a void segment centered around a Voronoi edge *E*_*i*_. (b) Visualization of nucleoid structure for three values of bending stiffness. The resulting radius of gyration *R*_*g*_*/R*, normalized on total cell radius, is shown below each image. Dashed blue lines enclose 95% of DNA, with values *R*_95_*/R* = 0.93, 0.94, and 0.864 from less to more stiff. (c) Distribution of pore sizes *d*_*i*_ (normalized on ribosome diameter) in the nucleoid [79]. (d) Average size ⟨*d*⟩ (r) of void segments located at distance *r* from the center of the cell, calculated using Equation (1). For both (c) and (d) we use data from 100 realizations.

**Figure 3(b)** illustrates visually how stiffness changes the pore mesoscale structure; the images indicate a progressively more heterogeneous structure as stiffness increases, with concentration of small pores / more DNA toward the center and larger pores / less DNA near the nucleoid’s edge, along with increasing DNA-free region. Simulation snapshots of the DNA backbone structure with inherent stiffness set at *σ*_*b*_ = 0, *σ*_*b*_ = 10, and *σ*_*b*_ = 50, respectively, are shown. Visual inspection indicates that inherent stiffness changes structure over several length scales, with apparent increase of persistence length with increased stiffness (see Figure S2 in *Supplemental Materials* for persistence length results). The corresponding DNA features thus change over shorter length scales with lower stiffness, but having flexible segments appears to promote spatially more-homogeneous structure. The stiffest nucleoid’s structure is visually more heterogeneous across the cell, with more compact structure near its center and a wider DNA-poor annulus near the membrane. The light blue dashed line in each snapshot encloses 95% of the DNA, illustrating this global compactness. To further quantify this global compactness, we also measured the radius of gyration — a commonly used metric in the literature — normalized on cell size, *R*_*g*_*/R*, for each stiffness condition. The result, shown below each snapshot, confirms that lower inherent DNA stiffness tends to expand the nucleoid region.

To quantify the stiffness-induced changes in nucleoid microstructure, we used the Voronoi volumes to measure pore-size distribution for 100 simulated nucleoids at each stiffness [**Figure 3(c)**]. Every pore *i* is scaled on ribosome size *d*_*i*_*/*2*a*_*rib*_ and is represented in the distribution. A single peak for each stiffness shows a dominant pore size, with weak skew toward larger pore size (we note that cells such as *E. coli*, which have a much larger genome, may form more complex mesoscale structures such as supercoils; these would produce at least on additional peak). For the stiffest nucleoid, more than half the pores are 75% the size of a ribosome (thus excluding them), and only 29% can accommodate a full ribosome. The peak height, location, and width all change appreciably for the less-stiff nucleoids, with only about half of pores excluding ribosome-sized biomolecules. While the simulation snapshots in (b) show a visual difference between the nucleoid structures for *σ*_*b*_ = 0 (many short-length scale bends, homogeneously distributed pore size) and *σ*_*b*_ = 10 (longer persistence length [see *Supplementary Material*], denser with smaller pores near the center), the pore size distributions in (c) are nearly identical for the soft and moderate stiffness, because these are averaged over the entire nucleoid.

But pore size *d*_*i*_ varies from center (*r* = 0) to membrane (*r* = *R*) more substantially as stiffness increases, as shown in **Figure 3(d)**. Recall that *d*_*i*_ is the diameter of a cylindrical segment of the network, which can be oriented in any direction. We construct a spherical surface *S*_*r*_ of radius *r* centered on the cell’s center, and identify any segment edges that intersect that surface. Any segment captured at *S*_*r*_ represents a volume or pore that a biomolecule can occupy. Systematically increasing *S*_*r*_ will capture the same segment until the increase in *S*_*r*_ is larger than the pore’s size — diameter or length, depending on orientation. Assuming that the orientational distribution of these segments (cylinders) is independent of distance from the cell’s center, this method appropriately captures how larger pores reside within the cell. The diameter of these segments is tabulated at each *r* and we plot, in **Figure 3(d)** the average size ⟨*d⟩* (*r*) of all voids located a distance *r* from the cell center using the expression

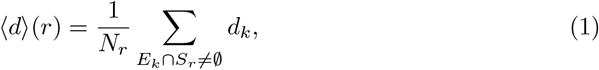

*i*.*e*., how void *sizes* are distributed spatially. Here, *N*_*r*_ is the number of edges (pores) intersecting a given spherical shell *S*_*r*_ and *d*_*k*_ is the size of the void identified by the edge *E*_*k*_ of the Voronoi network. **Figure 3(d)** shows that the change in bending stiffness from *σ*_*b*_ = 0 to *σ*_*b*_ = 10 drives down pore size near the center and drives pores size up near the edge of the nucleoid.

Evidently, the presence and concentration of DNA segments alone does not exclude ribosomes, but rather the mesh structure permitted by the stiffness of bonds. In fact, the stiffest nucleoid’s average pore size near the center is half the ribosome diameter, consistent with the exclusion of ribosomes observed in **Figure 2**. The two softer nucleoids’ central pore sizes are on average very close to ribosome size, potentially trapping ribosomes there [see *Supplementary material*].

### Gene accessibility, migration pathways and compactness

Knowledge of pore-size distribution is important to gain insight on nucleoid microstructure; however, this alone does not reveal dynamic accessibility of macromolecules moving within the nucleoid. Whether a diffusing RNAP, RNA, or other molecule can traverse through the entire nucleoid to freely access targeted genes depends on the *fully connected* pathways, or tortuous network, formed by continuously connected pores. A protein of a given size might be able to migrate partway from the edge of the nucleoid toward its center before becoming trapped, for example, whereas a smaller particle could freely explore the entire nucleoid. To explore this macromolecular accessibility, we used the Voronoi network constructed previously to identify connected pathways that can be traversed by biomolecules of finite size [79], by using a graph-traversing algorithm to identify connected components of a pruned network containing only pathways that permit passage of a biomolecule of size *a*. We call each accessible region defined by these fully-connected networks a *traversable region* [**see Figure 4(a)**]. Depending on the size of the biomolecule, there can be either a single, fully connected region, or multiple isolated regions. Note that these regions can either traverse the entirety (or the majority) of the nucleoid, allowing macromolecules such as transcription factors to access different genes, or, alternatively, trap large macromolecules. **Figure 4(a)** shows a zoomed-in image of one such region for a test particle with width 1.5 times the size of a ribosome. Any particle smaller than 1.5*a*_*rib*_ can move within this region successfully.

**Fig 4.**
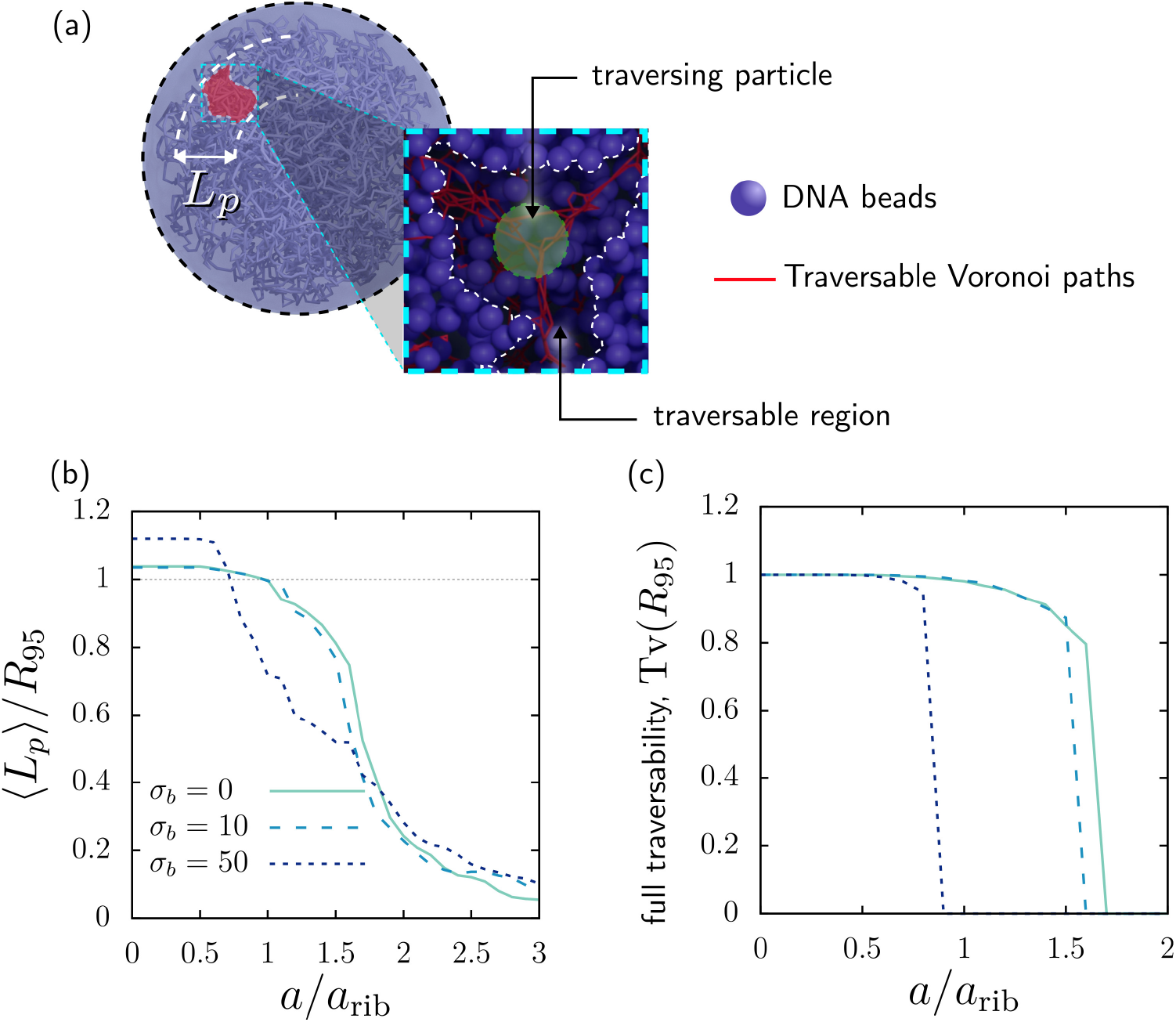
Traversability analysis of the nucleoid’s mesh structure, for three values of bending stiffness *σ*_*b*_. (a) Schematic of a biomolecule moving through a traversable region inside the nucleoid. The penetration length *L*_*p*_ (see annotation in (a)) quantifies the *radial* length of that region, *i*.*e*., the fraction of the distance from center to edge of the nucleoid. (b) Average penetration length *L*_*p*_ normalized by the radius of the region enclosing 95% of DNA content, *R*_95_, as a function of macromolecular size. The angle brackets represent an average over all isolated traversable regions [see *Methods* for details]. (c) Fraction Tv [Equations (18) and (19)] of all pathways that are fully traversable from center to edge (i.e., *L*_*p*_ ≥ *R*_95_), giving the likelihood that a molecule can traverse the entire nucleoid.

One insightful way to characterize these traversable regions is by measuring their penetration length *L*_*p*_, defined as the radial extent of motion of particles moving in such region [**see Figure 4(a)**]. For example, a penetration length *L*_*p*_ = 0.9*R* means that a particle in the outermost external portion of that region from the cell center, *R*_max_, can potentially move closer to to the center of the cell to a radial distance *R*_min_ = *R*_max_ ™0.9*R* from the center of the cell and vice versa. We carried out this analysis for a series of biomolecule sizes, ranging from zero to three times the size of a ribosome, 0 *≤ a ≤* 3*a*_*rib*_. **Figure 4(b)** presents the average penetration depth [see *Methods* for details of the averaging process] as a function of biomolecule size, showing that, on average, particles of size *a/a*_*rib*_ ⪅ 0.6 have some access from edge to center of the nucleoid; this penetration deepens for less-stiff nucleoids, *σ*_*b*_ = 0, 10. For all three values of stiffness studied, biomolecules must be smaller than 1.75*a*_rib_ to penetrate further than 40% into the nucleoid. As traversability also pertains to exiting the nucleoid from the center, values of ⟨*L*_*p*_*/R*⟩ ⪅ 0.1 indicate that biomolecules are either totally excluded from the nucleoid or totally trapped in a pore with no exit paths available.

The resulting traversable pathways can also be used to identify the likelihood of a biomolecule being able to fully penetrate the nucleoid, or, alternatively, being trapped deep within the nucleoid. This full-penetration probability depends on the number of fully traversable pathways, as well as their sizes, which can be quantified, for instance, by the number of Voronoi paths in a given region. **Figure 4(c)** presents the likelihood of a macromolecule to be in a fully traversable pathway (i.e., with ⟨*L*_*p*_*/R*_95_⟩ ≥ 1) as a function of its size. Similarly to Ryu and Zia [79] and other works in percolation phenomena [81], the increase in size leads to a geometric transition at a so-call *critical size* where particles are not longer able to fully traverse the nucleoid and, instead, get excluded or trapped. For *σ*_*b*_ = 0 and 10, these critical sizes are 1.6*a*_rib_ and 1.75*a*_rib_, respectively, indicating that most biomolecules the size of a ribosome are able to fully traverse the entire nucleoid. The weak decay from unity from 0.6 ≤ *a/a*_*rib*_ ⪅ 1.7 indicates some chance of such particles being trapped in an unconnected pore. Beyond each steep drop in full traversability, there are so few fully-connected pathways that such biomolecules will never fully traverse the nucleoid. Increasing nucleoid stiffness to *σ*_*b*_ = 50 cuts the critical size in half, to approximately 0.8*a*_rib_, indicating that no ribosome is physically able to fully traverse the nucleoid.

Overall, the physiological stiffness modeled here for Syn3A, *σ*_*b*_ = 10, with only entropic interactions and charge-neutral biomolecules — a simplification we remove in subsequent sections — provides abundant microstructure that is dynamically accessible to biomolecules of all sizes. As noted above, this inherent stiffness still compacts the nucleoid, leaving a DNA-free region near the membrane that is inconsistent with experimental observations of fully expanded nucleoid. We continue in the next sections increasing the biological conditions not represented thus far in the model that may further expand the nucleoid.

### Local HU-induced bends induce further nucleoid compaction

The results just presented indicate that a less-stiff nucleoid tends to be more expanded than a stiffer nucleoid, showing that inherent stiffness promotes DNA compaction, and ribosome exclusion is coupled to nucleoid inherent stiffness. However, the presence of the biomolecules within the nucleoid may produce entropic effects that enhance or compete with these enthalpic effects; we will examine these in subsequent sections below. Biomolecules such as nucleoid-associated proteins (NAPs) can also promote structural changes in the nucleoid by binding to DNA. In contrast to other bacteria, Syn3A only produces one type of nucleoid-associated protein: HU [82], which can not only make the nucleoid stiffer, but also produce local bends [13].

As explained in the Introduction, nucleoid-associated proteins (NAPs) such as H-NS are known to influence nucleoid compaction [17]. Similarly, we expect the HU proteins in Syn3A to also influence nucleoid compactness and ribosome distribution. Namely, the concentration of HU in Syn3A and other *Mycoplasmas* is notably lower than that of most bacteria [21], indicating that the paucity of these molecules may contribute to a lower degree of nucleoid compactness. In constrast to H-NS, which can create physical bridges between widely separated genes, HU proteins are known to increase nucleoid stiffness and/or create localized bends across a few DNA base pairs [**Figure 5(a)**], depending on their binding mode [13]. As shown in the previous section, an increase in DNA stiffness can result in a more compact nucleoid. In this section, we aim to understand to what extent the low abundance of HU, and the local bends generated by these proteins, contribute to the expanded nucleoid in Syn3A and other *Mycoplasmas*.

**Fig 5.**
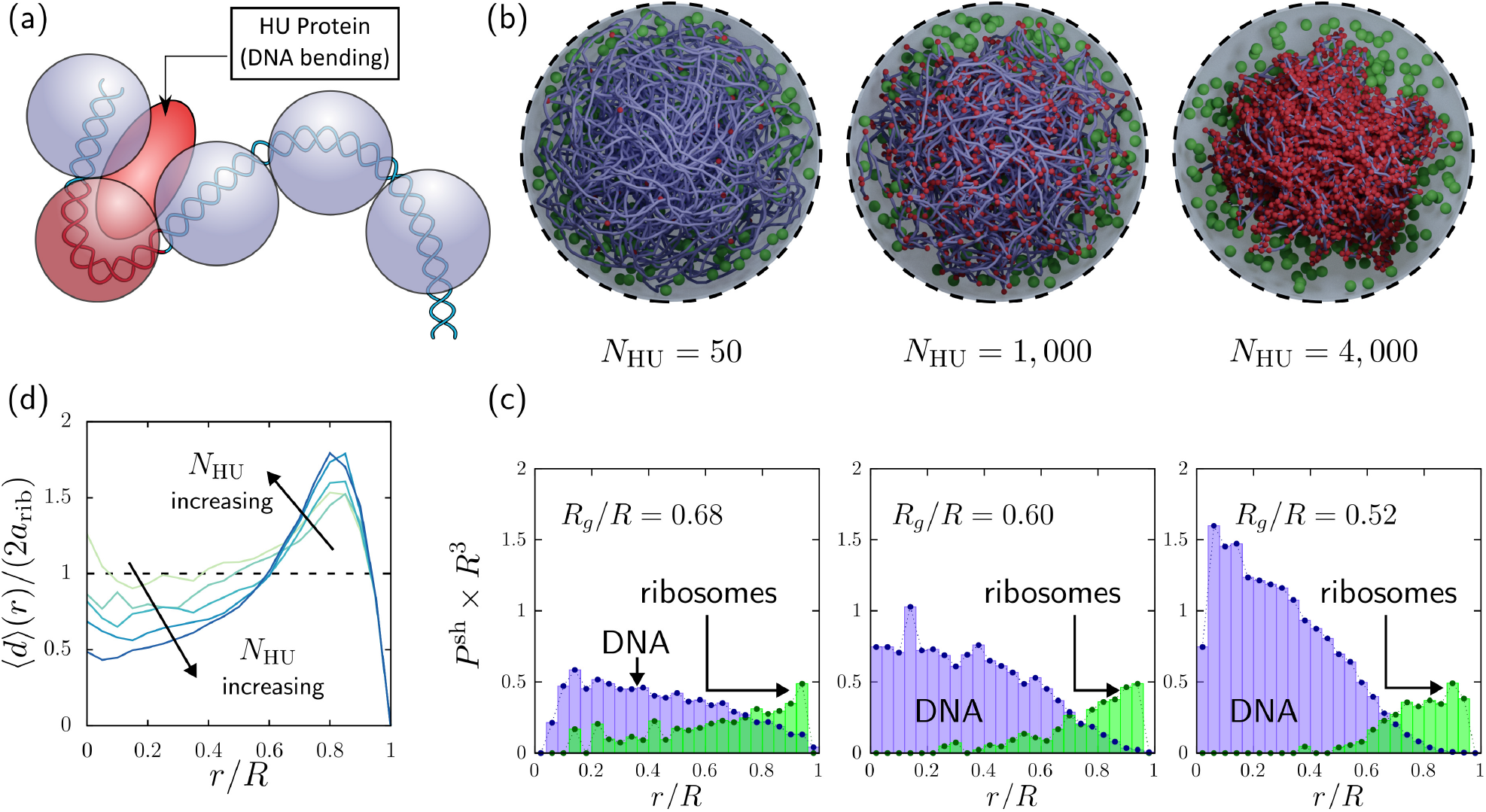
Influence of DNA-bending HU proteins on nucleoid compactness. Panel (a) illustrates schematically how an HU protein binds to DNA, creating a local bend with a local stiffness 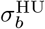 (in contrast to the inherent DNA bending stiffness *σ*_*b*_), and a local equilibrium angle *θ*_HU_ different from the typical *θ*_0_ = *π* used globally. To mimic how HU proteins bind to DNA independent of gene sequence, we induce bends in a random distribution throughout the genome (see *Methods*). Here we impose *σ*_*b*_ = *σ*_HU_ = 10, *θ*_HU=2*π/*3_, and simulate a systematic increase of HU concentration, *N*_*HU*_, shown in the figure. (c) Spatial distribution of DNA (purple) and ribosomes (green) from cell center to membrane, for the three values of *N*_*HU*_ shown. (d) Effect of an increasing number of HU proteins on the average size ⟨*d*⟩ (r) of void segments located at a distance *r* from the center of the cell, calculated using Equation (1).

To include the effects of the local bends induced by HU proteins into our model, we distribute local *bending defects* across the genome by locally changing the local equilibrium angle. The number of defects was chosen to represent a range of concentrations from physiologically accurate to high concentrations. The three simulation snapshots in **Figure 5(b)** show visually that, as HU concentration increases, the resulting local remodeling of the nucleoid produces global compaction and ribosome exclusion. Similar to the results in Figures 2 and 3, we quantify these effects by measuring the spatial distribution of DNA and ribosomes [**Figure 5(c)**], as well as the distribution of pore sizes within the nucleoid [**Figure 5(d)**]. This systematic nucleoid compaction with increasing HU, quantified by a decrease in radius of gyration, is similar to the effect observed in **Figure 2** for increased bending stiffness, confirming HU’s mechanistic action. Notably, the physiological concentration of HU proteins generates little additional compaction; substantial compaction occurs only at concentrations well above those reported in Syn3A (but more typical of other bacteria [21]). These observations, combined with the previous results for bending stiffness, support the prior hypothesis that paucity of HU proteins in Syn3A contributes to the nucleoid’s cell-spanning structure [74]. However, HU proteins are a small portion of the proteins inside Syn3A. What happens when we take into account the remaining native protein crowders?

### Cytoplasmic proteins entropically exclude ribosomes from the nucleoid

Up to this point, our model — similar to the model by [57] — did not include cytoplasmic proteins. This produced the leading order effect of lower cell packing fraction, but here we examine whether the presence of the secondary size population induces additional effects. Indeed, it is well known that crowding from cytoplasmic proteins drives nucleoid compaction [11]. We expect that a more accurate representation of physical packing in our model will further compact the nucleoid, actually worsening the agreement with experimental prediction of a cell-spanning nucleoid.

Building up on our baseline composition used in the previous sections, we can introduce proteins by using our swelling Monte Carlo algorithm [See *Methods* and the *Supplementary Material* for additional details]. The resulting macromolecular configurations are shown in **Figure 6**. For convenience, **Figure 6(a)** recapitulates our results for the simplified macromolecular composition, only containing DNA and a physiologically accurate number of ribosomes. Panel (a) shows the simulation snapshot and (b) shows the distribution from center to cell membrane of ribosomes and DNA. Next, we generate a more detailed cell with a physiologically accurate relative abundance of proteins, ribosomes, and genome constituents [**Figure 6(c,d)**] for Syn3A at growth rate 0.396 db/hr. The corresponding natural volume fraction is ϕ = 10.4% — an increase of 32% from the original model (ϕ = 7.9%). This more crowded, physiological composition of proteins, ribosomes and DNA markedly compacts the nucleoid and expels ribosomes out of the center of the cell toward the membrane [panel (d)]. The DNA-depleted annulus near the cell membrane is also widened. This degree of compaction and expulsion was also seen in **Figure 2** for *σ*_*b*_ = 50 (right-most figure), which had a very stiff nucleoid, showing that that protein crowders, despite contributing a relatively small portion of total volume fraction, has as strong an impact on nucleoid compaction as an extremely stiff nucleoid. Overall, our results confirm that the physiological abundance of proteins, often neglected in simulations, substantially influences nucleoid compaction.

**Fig 6.**
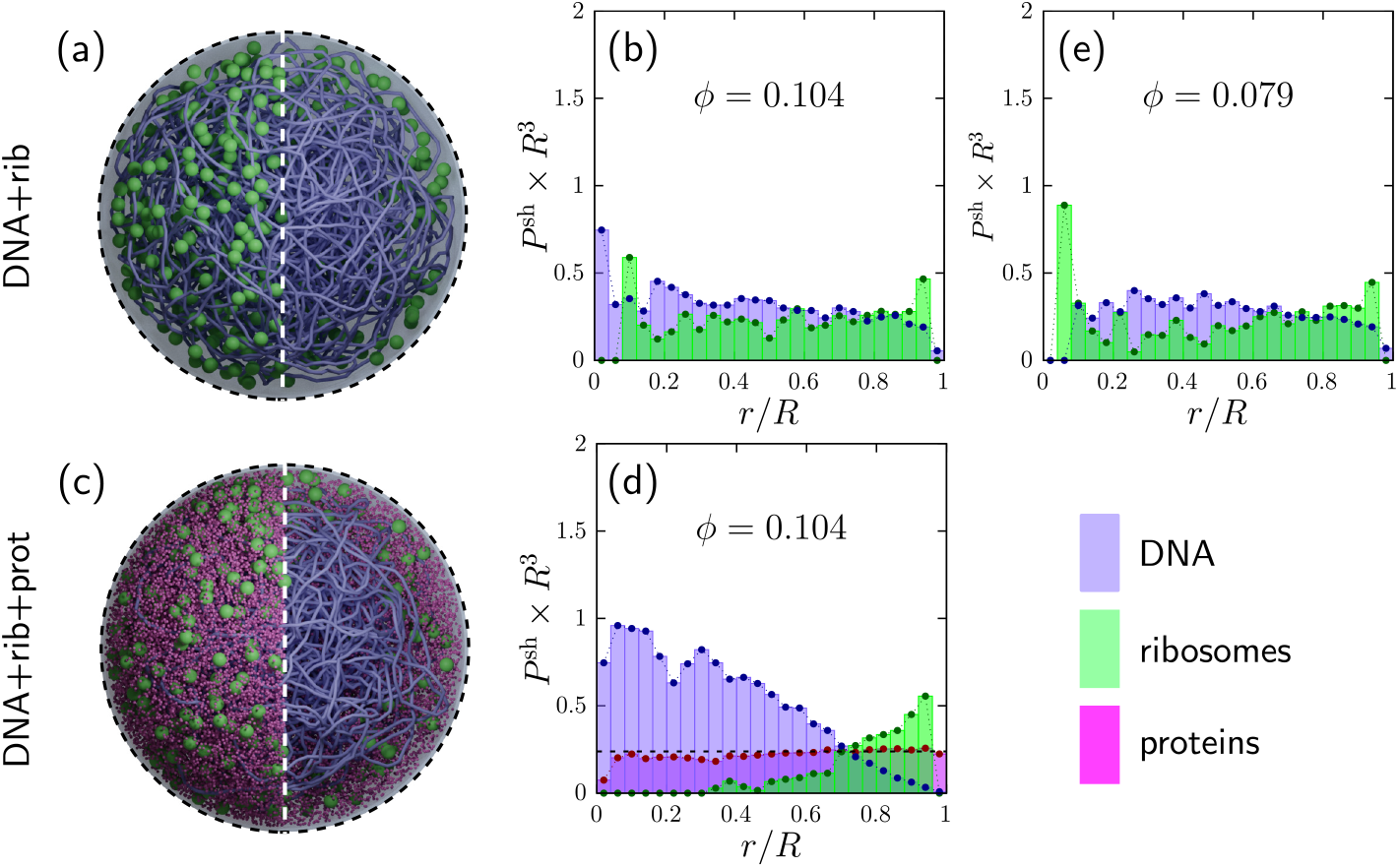
Dynamic simulations results for Syn3A model cell with nucleoid, ribosomes, and physiologically accurate abundance and packing of cytoplasmic proteins. Top row: no proteins. (a) Simulation snapshot; (b) Distribution of DNA (purple) and ribosomes (green), original model; (c) Same as (b) but at volume fraction corresponding to growth rate 0.396 db/hr. Bottom Row: with proteins. (d) Simulation snapshot; (e) complete set of ribosome, proteins, and DNA at growth rate 0.396 db/hr. Smaller biomolecules (proteins) drive more nucleoid compaction than larger biomolecules (ribosomes) at the same packing fraction, but a combination of both larger and smaller particles leads to preferential exclusion of larger biomolecules apparently not caused by nucleoid compaction.

The simplest mechanistic explanation for the increase in compaction is the increased volume fraction caused by the introduction of proteins into the model. To test this possibility, we simulated another version of our simplified composition (ribosome+DNA) and keeping the same volume fraction from panel (d) [**Figure 6(e)**]. Here, all proteins from (d) are replaced by ribosomes, which are five-fold larger than proteins. In agreement with the results in panel (b), the ribosomes extensively interpenetrate the nucleoid, forcing the nucleoid to expand to accommodate the larger ribosomes. This result suggests that there are other mechanisms at play other than the simple increase in volume fraction, such as a competition between entropic forces (not just proteins driving the nucleoid inward) that maximize entropy of DNA, ribosomes, and nucleoid. Mechanistically, we hypothesized that when both proteins and ribosomes are present, the combination maximizes entropy via proteins preferentially locating within nucleoid pores where their small size affords high vibrational entropy within pores, and ribosomes preferentially locate in the DNA-depleted periphery, where they gain more long-range (configurational) entropy [83]. This *combination* entropically promotes nucleoid compaction. Additional simulations show this to indeed be the case [see **Supplementary Figure S4**] and reveal an interplay with heterogeneous mesh size in the nucleoid: in more crowded conditions, proteins can maximize their entropy by moving to larger pores near the edges of the nucleoid, and by expanding the DNA-free surrounding region. The resulting osmotic pressure equilibrates by proteins further compacting the nucleoid until a balance is reached, where the nucleoid’s enthalpic energy then can reduce via compaction.

Overall, the loss in entropy for a more compact nucleoid is offset by the gained configurational entropy of ribosomes in the less-crowded periphery and maintained vibrational entropy of proteins within pores. This physiological composition of cytoplasm with simplified interactome promotes more compaction than measured in Syn3A, reinforcing the idea that other interactions, in addition to these entropic forces, underlie *Mycoplasmas*’ cell-spanning nucleoid. Indeed, it is well known that proteins, ribosomes, and DNA carry electrostatic charge. Next we examine how such charge influences spatial distribution of these biomolecules.

### Effect of electrostatic interactions in nucleoid compaction and organization of cytoplasm

To probe the effect of electrostatics, we endowed each biomolecule with surface-distributed electrostatic charge. All DNA and ribosomes in the model have uniformly distributed negative surface charge, with physiologically accurate values taken from literature. For protein surface charge, we determined the solvent-accessible residues of the folded protein structure and evaluated its dissociation constant to determine pH-dependent net charge. In **Figure S1(b)** (See *Supplementary Material*), we show the net charge and size of each of Syn3A’s cytoplasmic proteins. **Notably, 71 - 75% of Syn3A’s proteins are positively charged**, with 25 - 29% negatively charged (of these, a few only weakly so). **This is almost opposite to *E. coli*, whose proteome is about 40% positively charged**, and 60% negatively charged. These sizes and charges of ribosomes, DNA, and proteins is encoded into the interaction potential [see *Methods* and Supplementary Information]. The initial distribution of DNA, ribosomes, and proteins was first equilibrated without charge [**Figure 6(c,d)**]. The simulation then commenced, and particles immediately began interacting according to charge and entropic exclusion while Brownian motion and fluctuations proceeded (Equation (8)).

The simulation results are presented in **Figure 7**. A simulation snapshot in **Figure 7(a)** is split into two hemispheres, with some molecules in the right half made invisible to reveal the nucleoid’s shape. Aggregates and condensates have formed due to interactions between positively and negatively charged proteins and ribosomes. Several exploded views are shown around the main cell image to illustrate the range of composition and size of these condensates.

**Fig 7.**
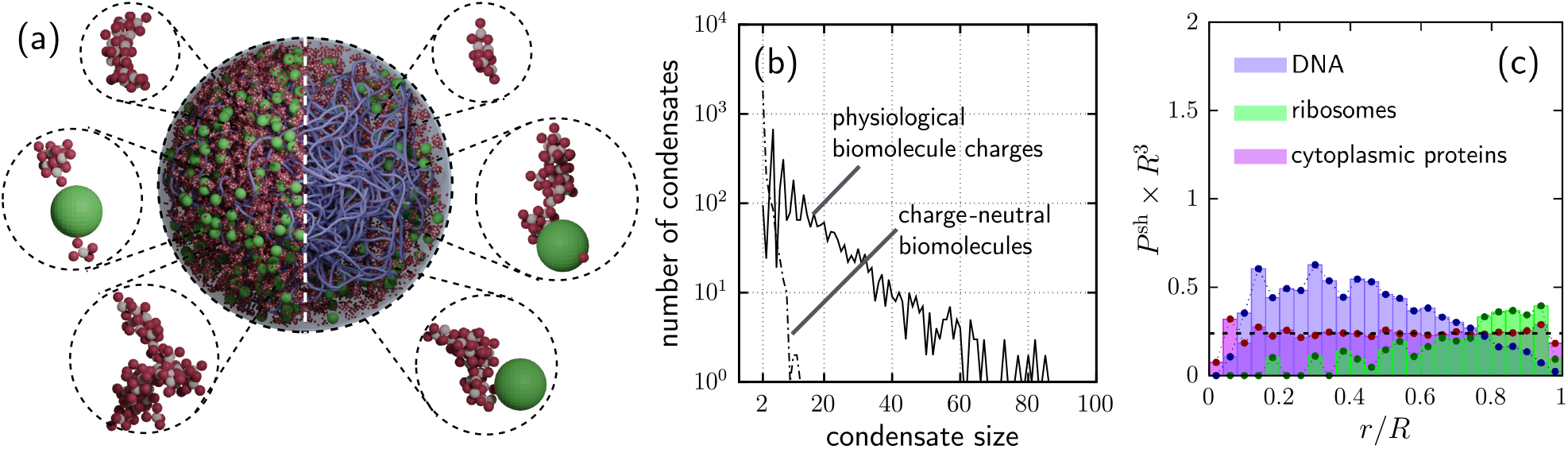
Influence of electrostatic interactions on nucleoid and cytoplasm organization in our Syn3A model cell. (a) Center image: equilibrated whole-cell simulation snapshot [DNA (purple), ribosomes (green), and proteins (positively charge are red, negatively charged are white)]. Six zoomed-in images surrounding the center image show protein condensates of various sizes, as indicated in the figure. (b) Condensate size distribution (in number of components) for charged molecules (solid curve) and neutral molecules (dot-dashed curve). Data for a single simulation. (c) Radial distribution of DNA (purple), ribosomes (green), and proteins (pink).

These sizes and compositions are analyzed in **Figure 7(b)** following the process described in *Methods* and in the Supplementary Information. The corresponding result for the uncharged system is also shown for comparison. Unsurprisingly, electrostatic interactions drive formation of small, medium, and large condensates. Some condensates comprise proteins only, while other involve proteins bound to ribosomes, an effect observed in other bacteria [7]. Interestingly, negatively-charged ribosomes are now present deep within the negatively-charged nucleoid. Analysis of these ribosomes reveals that they are part of condensates, coated with positively charged proteins. **Figure 7(c)** shows a substantial population of ribosomes within the nucleoid — agreeing with experimental evaluation of Syn3A and other *Mycoplasmas*. Evidently, “charge shielding” of the ribosome can drive their cell-wide distribution in Syn3A and other *Mycoplasmas*. These results are consistent with our recent study of *E. coli* [7].

**Figure 7(c)** further reveals that electrostatic interactions change the molecular distribution compared to the all-neutral biomolecules from **Figure 6(d)**. The key new result is that the electrostatic charges on proteins, DNA and ribosomes causes the nucleoid to expand throughout the cell. One key change is the reduction of DNA near the center: a shift of several thousand beads from the center re-distributes into the annulus near the membrane which, because it has significantly more volume, shows a graphically modest change. Concomitantly, ribosomes re-enter the nucleoid farther from the cell membrane. The resulting distribution better mimics the experimental results imported directly into a computational model [19, 57], as compared to models with no proteins or uncharged biomolecules.

## Discussion and final remarks

Nucleoid compaction in bacterial cells has been extensively attributed to supercoil condensation, NAP bridging, and cytoplasmic crowding [11, 12, 17, 26]. In many cells, such as *Escherichia coli*, these mechanisms condense the nucleoid to a central volume within the cell. In contrast, the nucleoid in many *Mycoplasmas*, including the synthetic minimal cell JCVI-Syn3A, derived from a *Mycoplasma*, span the entire cell. It has been speculated that lower concentration of nucleoid associated proteins (NAPs), as well as other proteins that induce supercoiling, underlies the expanded nucleoid in such bacteria. In parallel, the spatial distribution of ribosomes differs markedly between cells with compacted versus expanded nucleoids: highly concentrated outside the nucleoid versus a near-uniform distribution throughout the nucleoid (and cell), respectively. There is an important genotype-to-’physiotype’ implication of this physical distribution, namely, coupled transcription and translation [5], which in turn leads to different phenotypical outcomes. We coin the term ‘physiotype’ to capture how physical effects determine phenotypic outcomes.

Recent experimental studies of Syn3A reported this nearly uniform spatial distribution of ribosomes [19], leading to construction of a computational model with that ribosome distribution and a nucleoid self-assembled in the remaining space [57]. Syn3A has the simplest genome and proteome capable of life, and its NAPs are limited to a single type, the HU protein, making it an ideal system for studying the expanded-nucleoid physiotype. Indeed, recent discussions [74] surfaced a hypothesis that the paucity of HU proteins in Syn3A was the main reason for the expanded nucleoid configuration and uniform ribosome distribution. However, the only prior models of Syn3A included only DNA and ribosomes, which were all charge neutral [57, 65], providing a natural segue to a more refined model.

We built a computational model of Syn3A in a Brownian dynamics framework and reconstructed the ribosome distribution reported in the prior measurements and models [19, 57, 65]. We grew a self-assembled nucleoid via a swelling Monte Carlo algorithm [84, 85], initially maintaining the charge-neutral conditions previously modeled. However, we found that such conditions do *not* lead to homogeneous distribution of ribosomes. We resolved this conflict by recognizing that the prior models had recapitulated experimental distribution of ribosomes at their initial condition, but it turns out that this distribution is dynamically unstable, meaning that as soon as Brownian dynamics commence and molecules begin to diffuse, they redistribute into a heterogeneous configuration. This dynamical evolution of the system leads to an accumulation of ribosomes near the cell membrane that is inconsistent with experiments, further inspiring a more refined model.

We modeled inherent DNA stiffness and changes in nucleoid microstructure induced by HU proteins that bind to and form bends in the DNA, finding that physiologically accurate representation of these conditions again further compacted (rather than expanded) the nucleoid. Protein crowding amplified these effects: we added (charge neutral) cytoplasmic proteins to our model, with relative abundance and packing fraction taken from literature. We found that explicitly representing the physical presence of Syn3A’s cytoplasmic proteins — missing in prior models — actually exacerbates ribosome expulsion from and compaction of the nucleoid. That is, if prior models had included proteins and dynamics, compaction would have clearly emerged. We next endowed the proteome with physiological distribution of electrostatic charge (missing in prior models), which finally produced expansion of the nucleoid.

Overall, our results indicate that an interplay between inherent DNA chain stiffness, HU-induced bends, cytoplasmic crowding, and electrostatic interactions all play a role in the macromolecular distributions of DNA, ribosomes, and proteins in Syn3A. Increased nucleoid stiffness, HU concentration, and protein crowding all enhance nucleoid compaction, whereas electrostatic interactions are the only mechanism studied here that counteracts this compaction. Specifically, we find that the atypical charge distribution in Syn3A — in which the majority of proteins are positively charged — plays a role in the observed macromolecular organization by allowing more ribosomes to interpenetrate the nucleoid by shielding their negative charge [7].

Electrostatic interactions also drove condensate formation. Condensates are hypothesized to play an important role in bacterial metabolic processes such as gene regulation and metabolic compartmentalization. This physiotype behavior warrants further exploration; for example, our proteins’ surface charges were isotropic, and had a single value for all positively-charged proteins and a separate single value for negatively-charged proteins, and all with a single average size. Our model is capable of representing patchy distribution of surface charge, bumpy shape, and the physiological distribution of hydrodynamic size, all of which may yield interesting results, potentially revealing which parts of the proteome drive condensation, and genotype-physiotype-phenotype connections including stress adaptations. Electrostatic interactions may also involve membrane proteins related to cell division and other behaviors.

The nucleoid’s mesh structure also plays a crucial role in macromolecular separation and trapping, as demonstrated by our traversability analysis. We showed that morphological changes in DNA influence dynamic accessibility of macromolecules to certain portions of the nucleoid, such as accessibility to rrn operons or the replication origin. Recent studies speculate that such mesh structure fosters polysome formation deep within *E. coli*’s nucleoid [2, 80]. We expect that this tortuous-path behavior inside the nucleoid will also impact metabolic processes. For instance, even for a fixed DNA concentration (compaction), the difference in available paths can hinder accessibility of transcription factors to specific genes, as well as induce macromolecular entrapment and potentially promote RNAP foci, which will, in turn, induce changes in transcription rates.

The methods and frameworks developed in this paper can be further developed as noted above for proteomic detail, as well as molecular-scale explicit representation of processes such as mRNA translation [9, 10] and transcription, and used to study other bacterial systems.

## Methods

### Intermolecular interactions

Proteins, ribosomes, and DNA beads are represented in our model as interacting spheres suspended in a Newtonian cytosol. This coarse-graining approach is consistent with that taken in other recent approaches for whole-cell modeling [7, 57, 86]. Interactions between ribosomes and proteins, DNA beads, and other proteins, as well as protein/protein interactions include electrostatic attraction and repulsion, as well as hard-sphere exclusion. The latter, hard-sphere exclusion, is modeled via

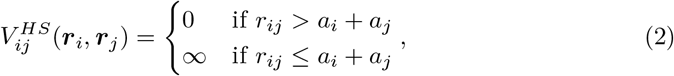

where *r*_*ij*_ = ‖***r***_*i*_ ***r***_*j*_‖ is the distance between the two beads *i* and *j* centered at ***r***_*i*_ and ***r***_*j*_ with radii *a*_*i*_ and *a*_*j*_. A similar interaction potential is also used to model interactions between macromolecules and the cell membrane. This potential prevents overlap between any contacting biomolecules in the simulation. For our *Brownian* simulations, the hard-sphere potential is replaced by a nearly-hard Morse potential, given by:

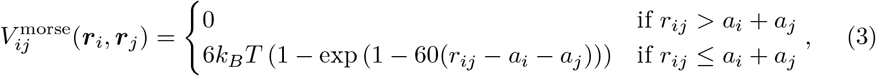

where *k*_*B*_ is Boltzmann’s constant and *T* is the absolute temperature. The potential in Equation (3) has been shown to recover hard-sphere behavior for volume fractions up to 50% [87, 88].

We modeled the circular bacterial chromosome as a closed-loop bead-spring polymer chain, consistent with previous DNA modeling approaches [76, 89]. The beads interact with each other via electrostatic attraction and repulsion, as well as hard-sphere exclusion, and also spring-like forces between base pairs and beads. The resistance to stretching and bending of the coarse-grained DNA chain is governed by spring and bending harmonic potentials, given, respectively, by

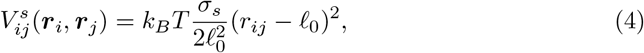

and

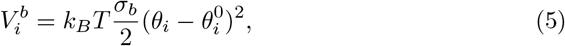

where ℓ_0_ is the equilibrium bond length between a pair of adjacent DNA beads. In Equation (5), *θ*_*i*_ is the angle formed by a triplet of beads consisting of the DNA bead *I* and its neighbors, and 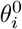 is the triplet’s equilibrium angle (see illustration, **Figure 8**). The dimensionless coefficients *σ*_*s*_ and *σ*_*b*_ are the *stretching stiffness* and *bending stiffness*, respectively. For “bare” DNA (i.e., in the absence of NAPs binding to DNA), the equilibrium angle 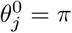 for all triplets throughout the entire chain. However, this equilibrium angle can be locally changed upon binding of bend-inducing HU proteins to the DNA [see **Figure 5** and corresponding discussion].

**Fig 8.**
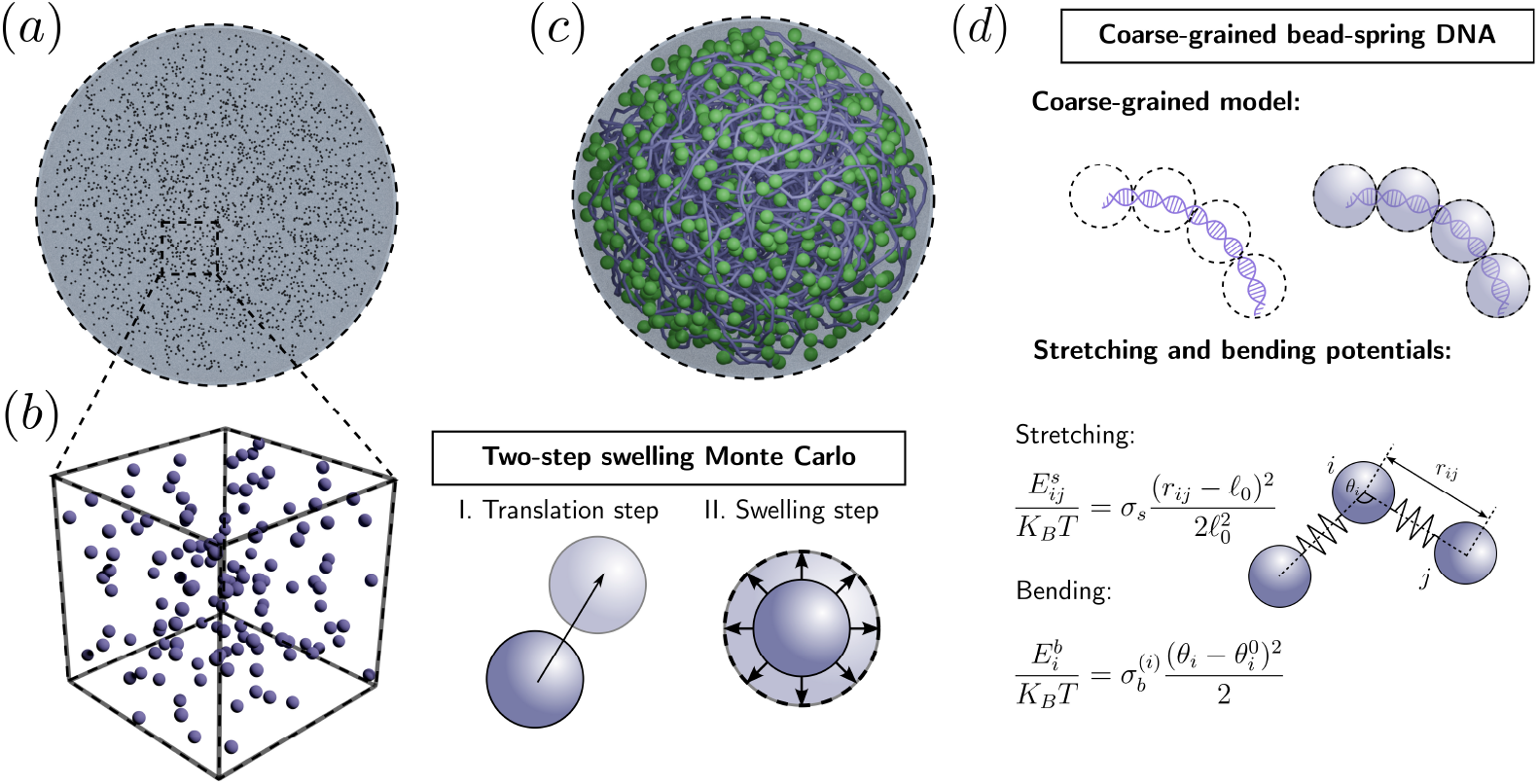
Illustration of the swelling Monte Carlo algorithm to generate cell configuration. (a) Initial positions of beads are distributed uniformly throughout the cell with assigned labels related to their respective types (including potentials and sizes). (b) Particles then undergo two Monte Carlo steps, translation and swelling. (c) Whole equilibrated cell. The DNA configuration is shown as a continuous connection of the backbone. (d) DNA beads (100 base pair each) are interconnected via combined bead-spring, bending, as shown in Eqs. (4) and (5).

Electrostatic interactions are represented via a screened electrostatic potential between two coarse-grained particles *i* and *j*, with nominal charges *q*_*i*_ and *q*_*j*_ [see *Supplementary Material* for details on determination of charges], given by a Debye-Huckel potential as:

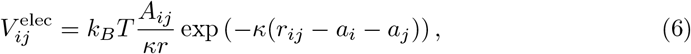

where *A*_*ij*_ is related to the net surface charges of particles *i* and *j* by [7, 90]

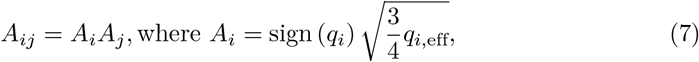

where 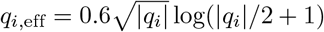 is the effective charge of the *i*-th particle and *κ*^*™*1^ is the Debye length.

### Swelling Monte Carlo algorithm

To generate the macromolecular distribution of ribosomes, proteins, and chromosome in Syn3A, we use an extension of the swelling Monte Carlo algorithm developed in [84, 91]. In contrast to some other approaches, such as the Koch-curve method used in [57] for chromosome generation, this method does not require any prior knowledge of experimentally measured positions of any macromolecules, allowing the physical interactions between particles to determine their location within the cell. In this approach, we first introduce all macromolecules inside the cell as points according to a uniform probability distribution and then perform translation and swelling Monte Carlo steps for each particles. An outline of the method is shown in **Figure 8**. Details about the method, including parameters and convergence criteria, can be found in the *Supplementary Material*.

### Langevin dynamics

For our mesoscale dynamic simulations, the motion of the coarse-grained macromolecules in the cell is governed by Newton’s second law, resulting in a Langevin equation:

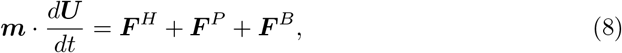

where ***m*** is a diagonal mass matrix, ***U*** is the velocity of all particles, ***F***^*P*^ are the forces due to interparticle and particle-membrane interactions, given by

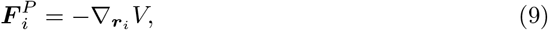

where

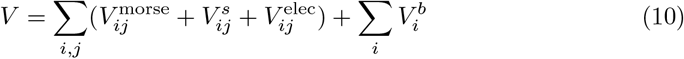

is the total potential energy of the system. Moreover, ***F***^*H*^ are the hydrodynamic forces, which, for spheres in the freely draining limit and in the absence of an external flow, are given by Stokes’ law ***F***^*H*^ = −6*πµ****a* · *U***, where ***a*** is a diagonal particle-radius matrix. Lastly, ***F***^*B*^ are the random forces and torques due to Brownian motion, given by ***F***^*B*^ = ***A* · Ψ**, where ***A*** is a positive-definite symmetric matrix and **Ψ**(*t*) is a white noise stochastic process, related to the hydrodynamic forces by the fluctuation-dissipation theorem.

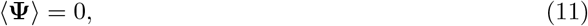

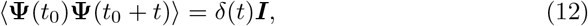

and

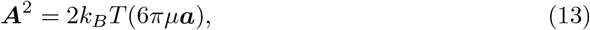

where ***I*** is the identity matrix, and *δ*(*t*) is the Dirac delta distribution. Equation (8) is numerically integrated using the langevin integrator in LAMMPS for a Stokes number *St* = 10^*™*4^, chosen to approach the overdamped limit of the Langevin equation. Similar to [7], the timestep was chosen to be 42 ps. Interactions between the macromolecules and the cell membrane are modeled similar to [7] by adding an additional force to Equation (8) that act inward in the radial direction given by

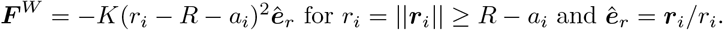

### Radial distribution function for single and multiple realizations

To quantify the distribution of DNA and ribosomes throughout the cell, we introduce reduced probability densities *P* (***r***) for each particle type. Due to radial symmetry, the true ensemble distribution only depends on the radial coordinate *r* = ‖ ***r*** ‖ and we write *P* (*r*). For a single numerical realization, the particle configuration is known and the probability density is given by

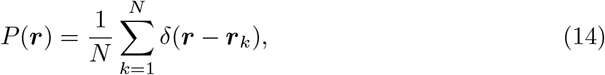

where *N* is the number of macromolecules of a given type and *δ* is the three-dimensional Dirac delta distribution. In this case, it is convenient to partition the cell domain into finite spherical shells of thickness *h* and calculate a shell-binned probability distribution, given by

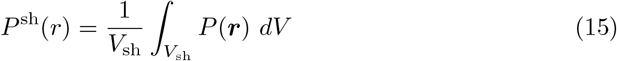

This approach, similar to previous works in confined systems [6, 92], quantifies the fraction of macromolecules whose centers are located in a spherical shell of radius *r* and thickness *h*, normalized by the shell volume. For our results, we use *h* = 2*a*_rib_. For a large number of particles *N*≫1 and small shell thickness *h*, this distribution approximates the real (ensemble) distribution function *P* (*r*).

For multiple runs, as we have many more available data points, the ensemble probability distribution *P* (*r*) can be better approximated by using spectral-decomposition methods [93]. To do so, we decompose *P* (*r*) in a series of orthonormal functions *ϕ*_*n*_ as *P* (*r*) = Σ_*n*_ *c*_*n*_*ϕ*_*n*_(*r*). The coefficients *c*_*n*_ can be calculated via orthogonal projection as:

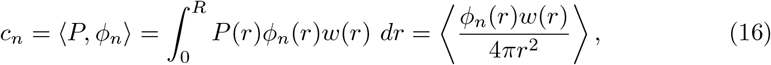

where *w*(*r*) is the inner product weight and the angle brackets denote an ensemble average. For numerical purposes, we truncate the Fourier series at a finite number of basis functions. To calculate the coefficients *c*_*n*_, we use a Monte Carlo quadrature to approximate the ensemble average in Equation (16) by an arithmetic mean over multiple particles at multiple runs, as in the methodology proposed in [93]. More specifically, we use the even <u>Cheby</u>shev polynomials as basis functions when expanding *P* (*r*), for which 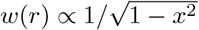.

### Calculation of average penetration length and traversability

The average penetration length ⟨*L*_*p*_⟩ shown in **Figure 4** is defined as

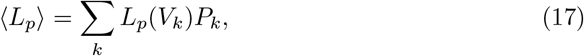

where the summation is performed over all connected components of the Voronoi network for a particle of size *a. P*_*k*_(*a*) is the probability of a particle of such size being in the traversable region defined by the *k*-th connected component of the Voronoi network *V*_*k*_(*a*). As the Voronoi network provides a good representation of the accessible regions for a given particle of size *a*, we make a simplifying assumption that there is an equal likelihood of a particle to be in any of the accessible Voronoi edges, meaning that *P*_*k*_ is given by 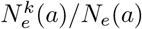, where 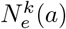 is the number of edges in *V*_*k*_(*a*) and *N*_*e*_(*a*) is the total number of Voronoi voids traversable by a particle of size *a*. Similarly, the traversability Tv(*α*), shown in **Figure 4(c)**, is defined as the probability of a particle being able to traverse a radial length *α* inside the cell. In **Figure 4(c)**, *α* = *R*_95_.

Mathematically, this is given by

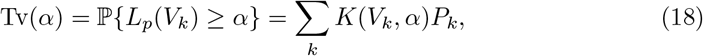

where the indicator function *K* is given by

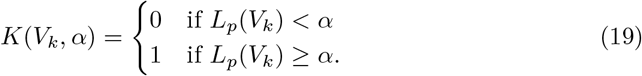

### Numerical determination of condensates

The determination of condensates in **Figure 7(a,b)** is performed by using a graph-traversal algorithm similar to the one used for our traversability analysis. Namely, we start by defining a connection network between beads, where edges are defined between particles *i* and *j* whose interparticle distance *r*_*ij*_ is smaller than a cutoff distance *c*_*ij*_ = (1.1)(*a*_*i*_ + *a*_*j*_). Identification of connected clusters in this network determines the agglomerates/condensates shown in **Figure 7**.

## Acknowledgments

The authors gratefully acknowledge funding support from the Alfred P. Sloan Foundation, Grant Number G-2022-19561. We acknowledge computational support of the University of Missouri Hellbender High Performance Computer Cluster. The authors thank Dr. John Glass (JCVI) for providing genomic data, the suggestion that HU proteins underlie Syn3A’s expanded nucleoid, and many useful conversations. The authors also thank the Build-A-Cell and SynCell communities for providing a robust community for exchange of ideas.

## Data availability

The experimental ribosome position data used to validate our method were obtained from a previously published work: Goodsell, D.S., Autin, L., “Integrative modeling of JCVI-Syn3A nucleoids with a modular approach” Current Research in Structural Biology, 7, 100121 (2024). The dataset is available from the original publication at https://doi.org/10.1016/j.crstbi.2023.100121. Post-processed data and code for analysis are available on Github https://github.com/gesseroure/Syn3Anucleoid.git. Dynamic simulations can be reproduced by the reader by utilizing the open-source platform LAMMPS [94]. Discussion regarding our particular modifications are available upon request.

## Author Contributions

**Conceptualization:** G.R., V.S.S., R.N.Z

**Data curation:** G.R., V.S.S.

**Funding acquisition:** R.N.Z

**Investigation:** G.R., V.S.S.

**Analysis:** G.R., V.S.S., R.N.Z

**Software:** G.R.

**Writing:** G.R., V.S.S., R.N.Z

