## Supplemental Information for "Abundant positively-charged proteins underlie JCVI-Syn3A’s expanded nucleoid and ribosome distribution"

### The swelling Monte Carlo algorithm for cell generation

To generate full macromolecular configurations without *a priori* knowledge of experimentally measured components of the cell, we expanded our swelling Monte Carlo algorithm, which is an extension of the algorithm developed previously for general packing and dynamics of random permeation networks in double emulsions [1,2]. We start by randomly distributing seed points inside the cell to generate a uniform distribution. Each point is then labeled to specify molecule type (here, DNA, ribosome, or protein), mesoscale structure (e.g., bonds or bending triples), electrostatic charge, and targeted size (radius $a_T$). For each Monte Carlo step, every particle undergoes two distinct steps: translation and swelling.

The translation step is sampled from a uniform distribution in a neighboring cube with sides $2h$, where $h$ is the amplitude of associated motion, set here to $h = 25$ Å to 50 Å depending on macromolecular composition. This translation step is subjected to a standard Metropolis-Hastings algorithm using the changes in the total system Hamiltonian, which includes entropic interactions (including boundary exclusion) and DNA bonding and bending potentials given by equations (4) and (5). A standard linked-cell-list algorithm is used to accelerate calculations. It is straightforward to incorporate the influence of electrostatic charge on the distribution resulting from this algorithm, as these potentials are well-defined in the framework, are attached to particle label vectors, and are currently used in our dynamic simulations.

In addition to translational displacement, each particle undergoes a swelling step. This swelling step is crucial for packing concentrated systems such as dense suspensions [3], because particles can adjust positions and organize, whereas placing full-size hard particles drives spurious overlaps that are computationally intensive to resolve, which becomes nearly impossible at high concentrations. Here, the initially point particles undergo a small change in radius at each Monte Carlo step at the swelling rate $\alpha_s = 0.1\%$ of the targeted final size $a_T$. In contrast to the energy contraints on the translation step, the swelling step is only subjected to a geometrical constraint: overlaps with other particles are disallowed:

$$a = \min\{a_0 + \alpha_s\, a_T, a_{cl}, a_T\},$$

where the clearance radius $a_{cl}$ avoids overlap. Two convergence criteria must be met: (i) all particles reach target size and (ii) convergence to the radius of gyration.

### Pipeline for extracting residue-level charge and size from PDB to colloids for mesoscale simulations

Each of the proteins in our model of JCVI-Syn3A is represented in our simulation as a physical object, with a mean surface charge and size. These proteins include RNAP, NAPs, and other well-characterized proteins. We built a database of all of these proteins, noting for each its relative abundance and its relative volume fraction at the single growth rate studied. There are at least three ways of characterizing size: two empirical formulations based on molecular weight [4,5] and on density [6], and a third based on the surface area of the folded protein, using data taken from the PDB. We used the third approach to estimate the size of the 419 proteins for which structure is reported for Syn3A, some of which are reported using homologs and others derived from AlphaFold predictions using sequence data. While our model is capable of handling an arbitrary particle size distribution, we opted to represent all proteins with a single size of 3 nm (compared to 13 nm for ribosomes and 6 nm for DNA beads), to simplify the analysis of entropic forces. To do so, we used our compiled data to compute the weighted average across Syn3A's proteome.

For surface charge of ribosomes and DNA, we used data available in the literature [7, 8]. For protein surface charge, we determined the solvent-accessible residues of the folded protein structure and evaluated its dissociation constant to determine the pH-dependent net charge using PROPKA3 [9]. We then used the Biopython `Seq Utils` library [10] to evaluate the molecular weight each proteins from its sequence. In **Figure S1(b)**, we show the net charge and size of each of Syn3A's cytoplasmic proteins, where the size is computed as noted above. **Notably, 71 - 75% of Syn3A's proteins are positively charged**, with 25 - 29% negatively charged (of these, a few only weakly so). **This is almost totally opposite to *E. coli*, whose proteome is about 40% positively charged**, and 60% negatively charged.

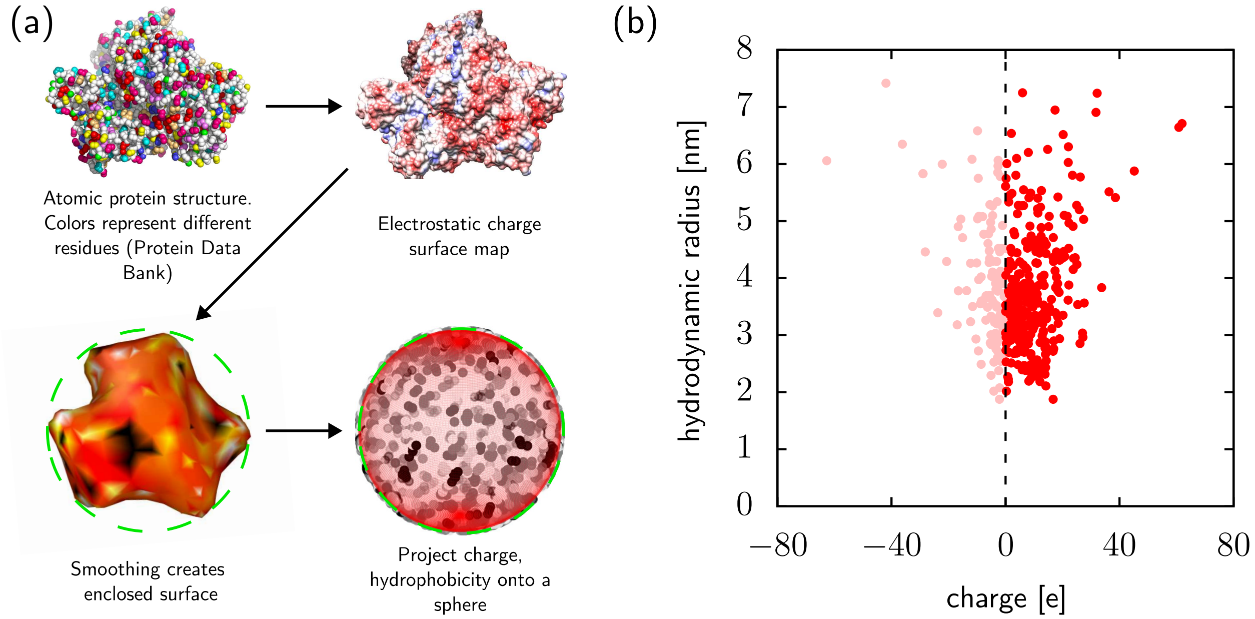


**Fig S1.** Determination of charge on and hydrodynamic size of JCVI-Syn3A's proteins. (a) Illustration of our computational pipeline for coarse-graining residue-level detail from Protein Data Bank structural data to a surface charge distribution and average. (b) Plot of the effective hydrodynamic size and net per-protein charge for each of Syn3A's proteins at pH= 7, obtained using PROPKA3 based on atomistic protein structures from PDB and AlphaFold. Plot showing the variation of net charge of cytoplasmic proteins with size.

### Persistence length of the DNA

One commonly-used metric for characterizing local remodeling of DNA is changes in the persistence length [11, 12]. Physically, the persistence length measures the length over which a flexible element such as a filament, polymer, or DNA strand remains 'straight' without bending or curving. Persistence length is related to mechanical rigidity or stiffness. In the case of DNA, such rigidity is set by inherent elastic properties, confinement, crowding, electrostatic interactions, and NAP interactions, among others. Persistence lengths in DNA can be obtained as the length over which two base pairs' orientation vectors (or two beads, in coarse-grained models such as ours) are correlated. This correlation is illustrated graphically in **Figure S2(a)**. The bond-correlation function is given by:

$$C(j) = \langle \boldsymbol{u}(i+j) \cdot \boldsymbol{u}(i) \rangle. \tag{1}$$

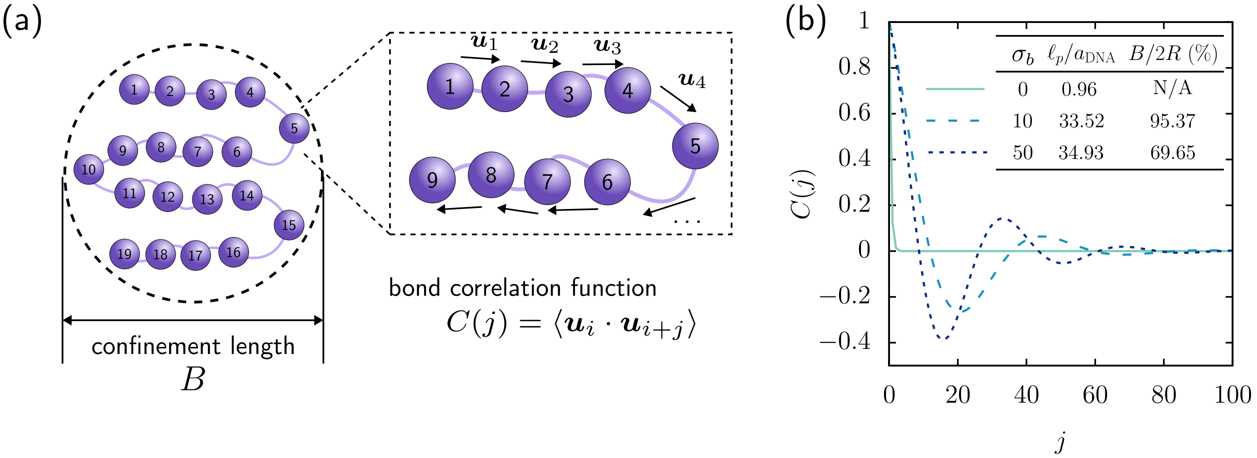


**Fig S2.** Bond correlation function and calculation of persistence lengths. (a) Illustration of the confinement length $B$ and bond orientation correlations that set the bond correlation function $C(j)$. The orientation vectors $\boldsymbol{u}_1$, $\boldsymbol{u}_2$, $\boldsymbol{u}_3$ and $\boldsymbol{u}_4$ demonstrate that the second and third bonds's orientations are closely correlated with each prior bond, but that the fourth bond starts to lose this correlation. This loss indicated the start of a bend, and the end of the persistence length. An undulation of these reversals forms a 'blob', measured as the confinement length $B$. (b) The bond correlation function was computed using Equation (1); a fit to that data using Equation (2) is shown in the figure. The persistence length (normalized on DNA bead size), $\ell_p/a_{DNA}$ and the confinement size (normalized on the cell diameter), $B/2R$, are shown for three values of DNA stiffness.

A useful phenomenological model of correlation length combines an exponential decay of memory, predicted by classical Flory theory [13, 14], which was later modified by Liu and Charkraborty to account for confinement [15]:

$$C(j) = \exp\left(-\frac{j\ell_0}{\ell_c}\right)\cos\left(\frac{2\pi j\ell_0}{B}\right). \tag{2}$$

We compute the correlation function $C(j)$ via Eq. 1. We obtain $\ell_0$ directly from simulations by measuring all bond lengths and computing the average value. Eq. 2 suggests that the peak-to-peak distance in the correlation functions for $\sigma_b = 10$ and $\sigma_b = 50$ correspond to the confinement length $B$. We measured this graphically (the peak-to-peak distance) and compared it with the value obtained by fitting the data to Eq. 2, which obtaining good agreement. This fitting process also yields the persistence length $\ell_p$. The amplitude decay from the first to the second peak is exponential, with decay strength corresponding to the persistence length $\ell_p$ (an exponential decay in memory of the previous orientation). The resulting fits, as well as the values of $\ell_p$ and $B$ for each stiffness value, are shown in **Figure S2(b)**.

For $\sigma_b = 0$, the amplitude decay $\ell_p$ is smaller than the size of a DNA bead. Recalling **Figure 3(b)** from the main manuscript, this nearly absent correlation is indicated by the sharp bends and short segments between bends that wipe out correlation between DNA beads. In contrast, higher inherent stiffness ($\sigma_b = 10$ and 50) drives fewer bends, as reflected in the larger values of $\ell_c$ and $B$. The confinement length, or "blob" size, suggests that as stiffness increases, there are discernible secondary structural length scales — effective stiffness is increased because there are reductions in conformational space induced by periodic repetition of correlation. Interestingly, this confinement length scale $B$ is remarkably representative of the nucleoid sizes observed in **Figure 3**.

### Voronoi and pore analysis

As discussed in the Results section of the main manuscript, we characterized the porous microstructure of the nucleoid via a Voronoi-based analysis following the approach by [16], with modifications that account for membrane confinement. Voronoi tesselation revealed pores of a range of sizes $d_i$ in the void regions excluded

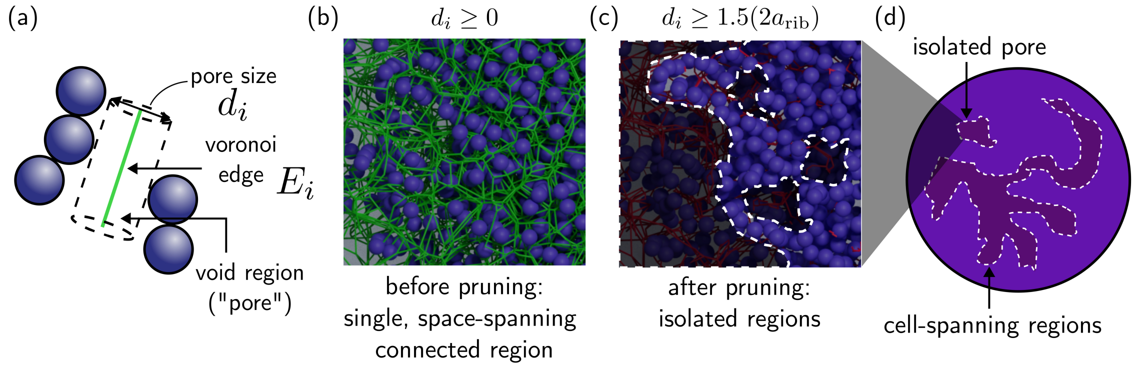


**Fig S3.** Pruning the Voronoi network creates isolated pores that can trap biomolecules.

by DNA and confinement. We omitted Voronoi edges near the membrane that produce spuriously high data for voids of nearly-null size. Here we provide additional information regarding how this analysis reveals pore structure and pathways.

**Figure S3(a)** again illustrates that a cylinder of diameter $d_i$ centered on a Voronoi edge 'just' fits within pore $i$. This diameter represents the maximum size of a biomolecule that can pass through that segment of a pathway. Panel (b) in the figure shows a section of the nucleoid model with the full Voronoi network, i.e., $d_i = 0$. In image (c), we have pruned the network to permit only particles of size $1.5a_{rib}$ or smaller. The image shows that not only are there fewer traversable pathways, but also, pruning automatically cuts off some pathways into isolated pores. Figure (d) shows one example of a fully-connected pathway across the nucleoid and one isolated pore.

### Accessible pathways inside the nucleoid

In the main manuscript, we analyzed the porous microstructure of the nucleoid using an extension of the methodology used by Ryu and Zia [16]. Here, we include additional visualizations of Voronoi pathways that illustrate how traversability of the nucleoid depends on the size of migrating particles and on inherent DNA stiffness. This visualization reveals how the pruning process eliminates impassable pathways.

When performing the Voronoi analysis, we associate each edge of the Voronoi network to a cylindrical void region through which particles smaller the cylinder's radius can traverse. As particle size increases, the number of fully traversable Voronoi pathways decreases (cf. **Figure 3**). To help visualize these traversable pathways, and how they change with bending stiffness, we took simulation snapshots near the central region of the cell. These snapshots are presented in **Figure S4** for three values of bending stiffness and four values of particle size. The image shows a slice through the center of the region to expose the pathways (centerlines shown in red).

As expected from the quantitative results in **Figure 3(d)**, for all values of stiffness, as the size of a migrating particle increases, the number of traversable pathways decreases. For $\sigma_b = 10$, and $a = a_{\mathrm{rib}}$, disconnects emerge, representing pores that may trap biomolecules near the nucleoid center. This trapping is quantified by a spike in ribosome concentration near the cell's center [e.g., Figures 1 and 6 in the main manuscript]. For bending stiffness $\sigma_b = 50$, biomolecules of size $0.75a_{\mathrm{rib}}$ can be trapped in small pores within the nucleoid. There are almost no pores that accommodate ribosome-sized particles or larger.

### Impact of crowder size on nucleoid compaction

In the Discussion section of the manuscript, we showed that the introduction of proteins into a simplified model Syn3A cell (DNA and ribosomes only), interacting only via entropic exclusion, expels ribosomes from the nucleoid, which is also compacted [see **Figure 6** and associated discussion]. This effect is not simply the result of coarse volume fraction, which we showed by instantiating instead an equivalent increase in volume fraction via added ribosomes, which are five times larger than the proteins modeled. This size-dependent effect of crowders arises from size-based segregation, where entropy is maximized by pushing the larger particles out to a DNA-free region, and proteins are able to maximize short-range entropy within the nucleoid's pores. Here we present results where we compare the relative impacts of size distribution and volume composition, holding total volume fraction constant, shown in **Figure S5**. Panels (a)(i), (b)(i), and (b)(ii) were previously presented in the main manuscript (**Figure 6**). Above, (a)(i) represents the physiological composition of Syn3A, (b)(i) a simplified composition (DNA + ribosomes) with same volume fraction, and (b)(ii) DNA with proteins only (no ribosomes). Panel b(iii) has the same amount of DNA as b(i) and b(ii). The bottom row shows our model cell with only DNA and proteins, with c(i) at the same volume fraction as (a)(i) and (b)(i), allowing the first column to compare the impact of proteins at the physiologically accurate volume fraction of Syn3A's DNA, ribosomes, and proteins; and (c)(ii) with the same volume fraction as (b)(ii) for the same comparison and, finally, (c)(iii), which has the physiologically accurate number of proteins and relative abundance to DNA.

In the main manuscript, **Figure 6** indicated that when both proteins and ribosomes are present, the combination maximizes entropy via proteins preferentially locating within nucleoid pores where their small size affords high vibrational entropy within pores, and ribosomes preferentially locate in the DNA-depleted periphery, where they gain more long-range (configurational) entropy [17]. This *combination* entropically

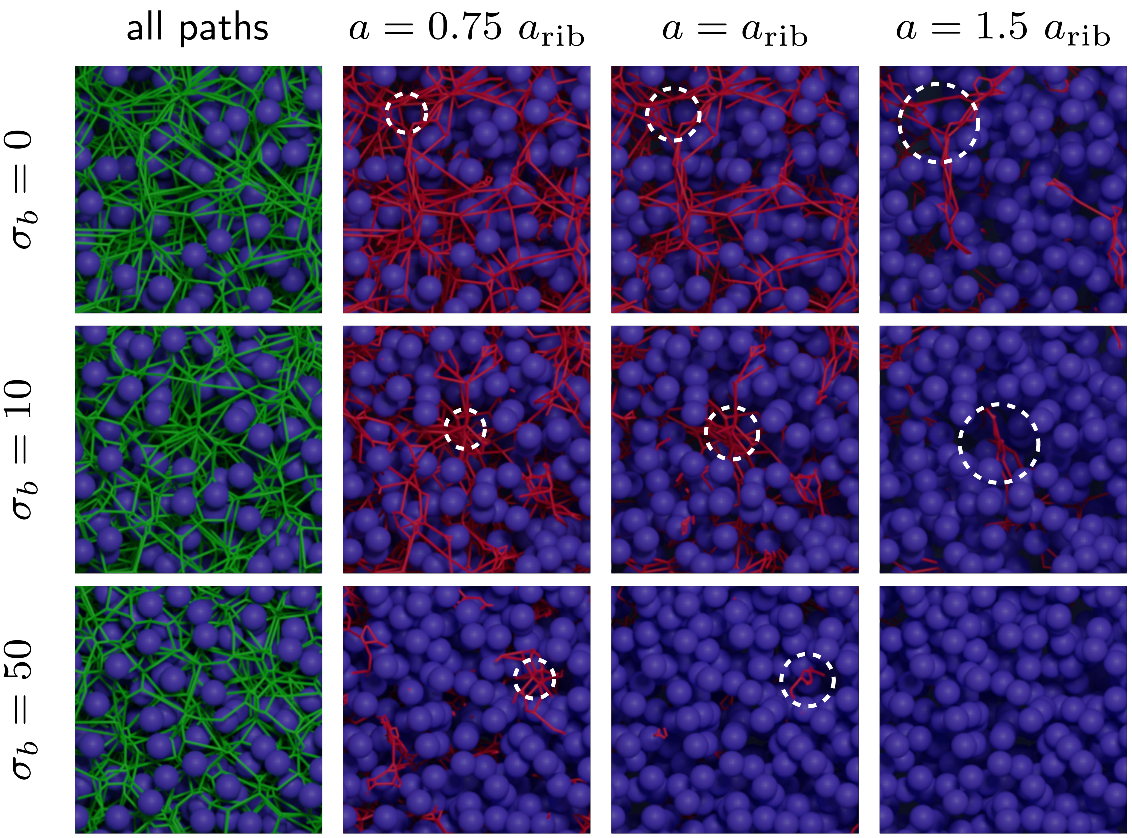


**Fig S4.** Illustration of how DNA stiffness changes nucleoid microstructure, pore sizes, and connectivity of the mesh network. Each row corresponds to a certain DNA stiffness, as shown. Vertical columns illustrate all paths traversable by a particle of maximum size shown at top. Green edges show the entire Voronoi network, where cylindrical pathways around each edge can be connected from pore to pore. For the first column, infinitesimally thin cylinders connect the greatest number of interconnected pathways, permitting passage of infinitesimally small particles. Finite-size particles require finite-width passageways (centerlines shown in red). Larger particles require wider pathways, which prunes the network to fewer available fully-traversing pathways.

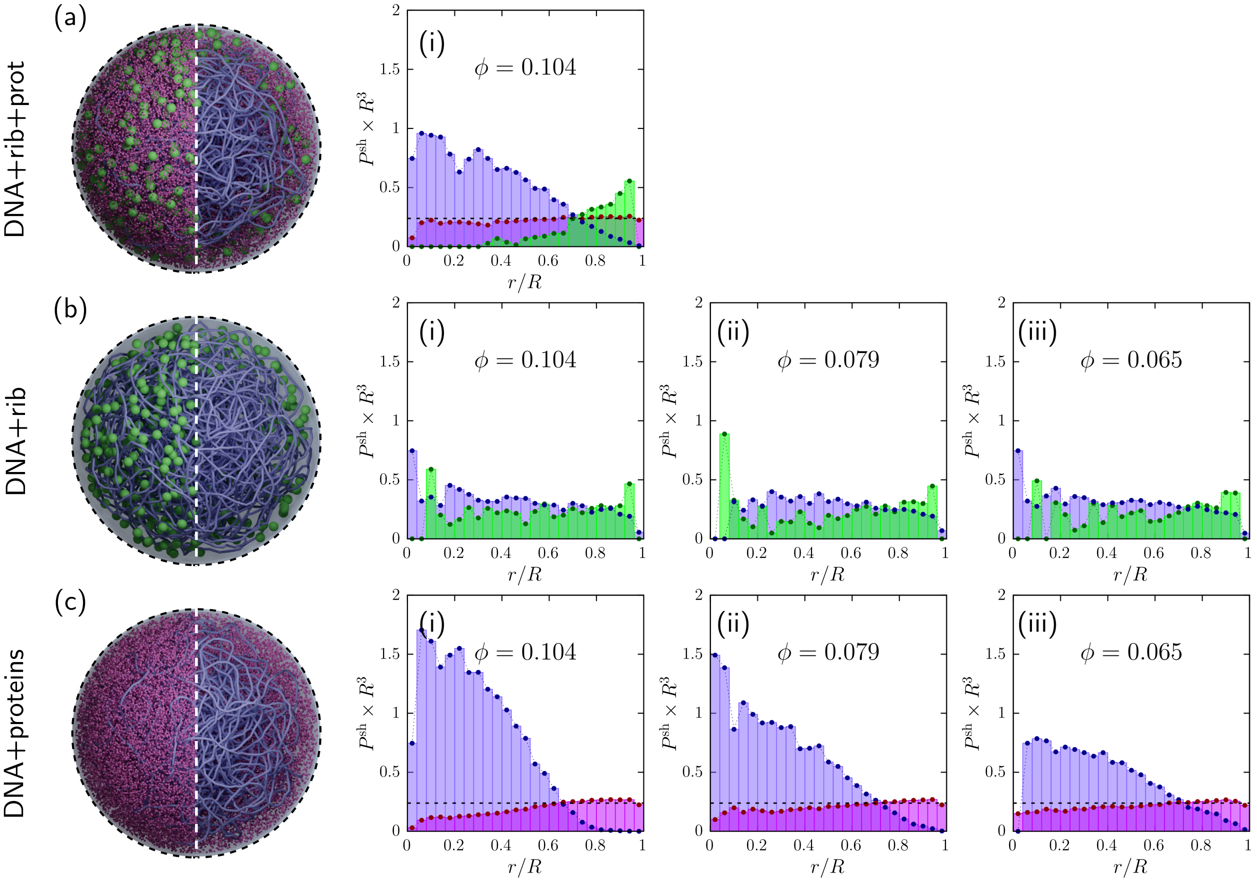


**Fig S5.** Effect of size composition and overall crowding (volume fraction, $\phi$) on distribution of DNA, ribosomes, and cytoplasmic proteins in our computationally modeled Syn3A cell. All biomolecules in this set of simulations are charge neutral. Size distributions obtained from PDB and our coarse-graining pipeline as discussed above. Each row focuses on a distinct molecular composition, with perturbation of volume fraction within each row: (a) DNA, ribosomes, and proteins; (b) DNA and ribosomes only; (c) DNA and proteins only.

promotes nucleoid compaction. To further understand how spatial distribution of DNA, ribosomes, and proteins is influenced by crowder *size*, we compare here how spatial distribution changes upon increased volume fraction induced by adding (b) only larger particles (ribosomes) and (c) only smaller particles (proteins). Comparison of the distributions in (b)(i,ii,iii) shows that increasing total cell crowding by adding more and more large particles (while holding DNA concentration fixed) induces almost no change in molecular distribution. In contrast, comparing (c)(i,ii,iii) show a marked influence of increased crowding on spatial distribution of molecules — where crowding was increased by adding more and more *small* particles (proteins). The nucleoid is up to 75% more compacted by this effect. These results indicate that ribosome exclusion and nucleoid compaction observed in (a)(i) are not caused solely by the increase in volume fraction, and that a competition between long-range (configurational) entropy of small particles throughout the cell versus short-range (vibrational) entropy within pores exerts a strong influence on nucleoid compaction. Proteins are able to maximize their entropy in more crowded conditions by moving to the larger pores in the outer region of the nucleoid, as well as by expanding the DNA-free surrounding region. The resulting osmotic pressure equilibrates by proteins further compacting the nucleoid until a balance is reached, where the nucleoid's enthalpic energy then can reduce via compaction.
